# Nascent transcription and the associated *cis*-regulatory landscape in rice

**DOI:** 10.1101/2022.07.06.498888

**Authors:** Jae Young Choi, Adrian E. Platts, Aurore Johary, Michael D. Purugganan, Zoé Joly-Lopez

**Author notes:** Corresponding authors (ZJL), (MDP) Coauthor email list (JYC), (AEP), (AJ).

## Abstract

**Background:** Plant genomes encode transcripts that require spatio-temporal regulation for proper cellular function, and a large fraction of the regulators can be found in intergenic regions. In animals, distal intergenic regions described as enhancer regions are actively transcribed as enhancer RNAs (eRNAs); the existence of eRNAs in plants has only been fairly recently documented. In this study, we evaluated with high sensitivity the synthesis of eRNAs that arise at genomic elements both distal and proximal to genes by combining PRO-seq with chromatin accessibility, histone modification, and methylation profiles in rice.

**Results:** We found that regions defined as transcribed intergenic regions are widespread in the rice genome, and many likely harbor transcribed regulatory elements. In addition to displaying evidence of selective constraint, the presence of these transcribed regulatory elements are correlated with an increase in nearby gene expression. We further identified molecular interactions between genic regions and intergenic transcribed regulatory elements using 3D chromosomal contact data, and found that these interactions were both associated with eQTLs as well as promoting transcription. We also compared the profile of accessible chromatin regions to our identified transcribed regulatory elements, and found less overlap than expected. Finally, we also observed that transcribed intergenic regions that overlapped partially or entirely with repetitive elements had a propensity to be enriched for cytosine methylation, and were likely involved in TE silencing rather than promoting gene transcription.

**Conclusion:** The characterization of eRNAs in the rice genome reveals that many share features of enhancers and are associated with transcription regulation, which could make them compelling candidate enhancer elements.

## Background

The spatio-temporal regulation of gene expression is essential for coordinating development and adaptive responses to environmental change. Transcriptional regulatory DNA sequences encode information that leads to the recruitment of transcription factors (TFs) in a DNA sequence-dependent manner, allowing the control of the location and rates of chromatin decompaction, transcription initiation, and finally the release of RNA polymerase II (RNAPII) from pausing to productive elongation[1, 2]. RNAPII can be recruited by regulatory elements both proximal and distal from the gene(s) they regulate. In animals, an important feature of many (if not all) active *cis*-regulatory elements (CREs), like enhancers, is the production of nascent transcripts by enhancers[3, 4].

Enhancer RNAs (eRNAs) have been generally defined as bidirectional, largely unspliced and unpolyadenylated transcripts originating from putative enhancers, with lengths predicted to be less than 150 nucleotides (although some can be up to 2 kb long) [5, 6]. In animals, eRNAs are predominantly localized in the nucleus and chromatin-bound fractions [7, 8], and are considered a hallmark of active enhancers and a proxy in predicting the spatio-temporal activity of active CREs. Enhancer transcription is important during gene expression. For example, it may maintain chromatin accessibility to enable the binding of TFs and cofactors [9, 10], it stimulates the catalytic activity of chromatin remodelers like histone acetyltransferases, it regulates the occupancy of TFs and coactivators [11, 12], and it promotes the pause release of RNA polymerase II to productive elongation [13, 14].

The instability of eRNAs, however, makes them difficult to detect with steady-state RNA-sequencing data, and their characterization depends on nascent RNA sequencing technologies that enable the measurement of transient RNA transcription at multiple stages and on a genome-wide scale. These methods include global run-on sequencing (GRO-seq) [15] and precision nuclear run-on sequencing (PRO-seq) [16]. These approaches have also successfully identified a diversity of RNA species, including long non-coding genes, upstream antisense RNAs, and eRNAs [17].

In plants, our understanding of eRNAs is still limited. There have been reports of eRNA transcription in maize and cassava [18], and in rice (*Oryza sativa*) [19], but they appear to be rare in other genomes such as Arabidopsis [20]. In maize and cassava, the intergenic regulatory elements from PRO-seq data appear to be enriched for expression quantitative trait loci (eQTLs) identified in kernels compared to a set of random intergenic regions and showed low levels of conservation, a pattern suggesting that these sequences evolve rapidly [18]. In rice, intergenic regions enriched for eRNA signatures had a marked enrichment for open chromatin, a generally asymmetric enrichment for H3K27ac histone modification, and weak but significant positive correlation with nearby gene expression. Similarly to maize and cassava, these sequences had low interspecies conservation but evidence of greater than two-fold excess of nucleotide sites under weak negative selection within *O. sativa* populations [19]. Evidence of recent selection on these sequences suggests the recent emergence of eRNA-producing intergenic regions within *O. sativa*, consistent with transcribed regulatory elements often being species specific [19, 21]. Understanding the relationship between eRNAs and the broader class of transcribed regulatory elements in plants is an important challenge and may in part be aided by methods that associate specific genetic variants with agronomic traits. These often indicate that these variants are located in noncoding sequences and likely functional in gene regulation [22, 23].

In plants, enhancer regions described so far appear to display specific characteristics. These include the presence of transcription factor (TF) binding motifs, chromatin accessibility, specific histone modifications, low DNA methylation, and evidence of physical interactions with their target genes that may be transient or stable [24, 25]. The benefits of considering several of these signatures in parallel has increased our ability to differentiate enhancers from other types of CREs (e.g. silencers, insulators, TATA-box, etc.), as well as other types of regions like promoters, transcription start sites (TSSs) and coding regions [19,24,26–31]. In addition, coupling these signatures with massively parallel reporter assays have greatly contributed to further functional characterization of CREs [26,32,33]. For example, a large proportion of the DNA in regions with active CREs appears more hypomethylated compared to other intergenic regions [28,34–37]. The presence of histone modifications also signal the presence of either active CREs (e.g. H3K27ac, H3K18ac) [38, 39], CREs with paused polymerases (enriched with H3K27me3 and low levels of an active histone mark such as histone acetylation) [26,40,41], or repressed CREs (low chromatin accessibility and and H3K27me3 levels) [26,42,43].

Despite the analysis of the various genomic and epigenomic features associated with plant enhancers, the presence of eRNAs are generally not considered. In this study, we used a combination of PRO-seq, complementary functional genomic datasets (DNA methylation profiles, histone modification profiles, transcriptomic profiles, chromatin accessibility), and 3D genome architecture to explore eRNAs and their genomic contexts in the Asian rice genome (*Oryza sativa*). We found that some intergenic eRNAs can be related to transposable element (TE) silencing but many share features of enhancers that suggest they harbor transcribed regulatory elements and are associated with transcription regulation and *cis*-expression quantitative trait loci (e-QTLs). This work continues efforts to understand important types of noncoding elements and is driven by the need to improve crop species by modulating gene expression to enhance plant system resilience [44].

## Results and Discussion

### Profile of nascent transcription in the rice genome

The goal of the study was to investigate the association between intergenic transcription and putative regulatory sequences in the context of DNA sequence composition, chromosomal environment, and evolutionary constraint. To examine genome-wide nascent transcription, particularly intergenic transcription including eRNAs, we generated PRO-seq data from rice leaf tissues of the *O. sativa* japonica cultivar Azucena grown under optimal conditions. We combined the newly generated PRO-seq data with previously published PRO-seq data that was also generated from plants grown under the same condition [19]. These PRO-seq datasets were combined and consisted of 17,519,424 reads that were aligned to the Azucena reference genome [45].

Because of the bi-directional nature of eRNAs [46, 47] and based on previous nascent transcription studies in animals [e.g. [48–52]], we used bi-directional transcription activity as a first criterion for identifying candidate transcribed regulatory elements in the genome [7, 53]. To do so, we used the machine learning algorithm dREG (discriminative Regulatory Element detection) that used mapped reads from our PRO-seq experiments (positive and negative strands) and used a support vector regression to recognize the characteristic pattern of divergent transcription at active transcribed regulatory elements (promoters, enhancers, and insulators) [50, 54]. Genomic regions with significant bidirectional transcription (herein referred to as dREG peaks) showed an enrichment of PRO-seq reads in both DNA strands, and highly active regions were assigned a high dREG score (>1.0; Fig. 1a). In total we detected 69,898 dREG peaks across the genome, with dREG scores ranging from 0.33 and 1.56 (Fig. 1a; Additional file 1: Figure S1). The genome-wide median dREG score was 0.591 and the median size of a dREG peak was 390 bps (Additional file 1: Figure S1).

**Figure 1.**
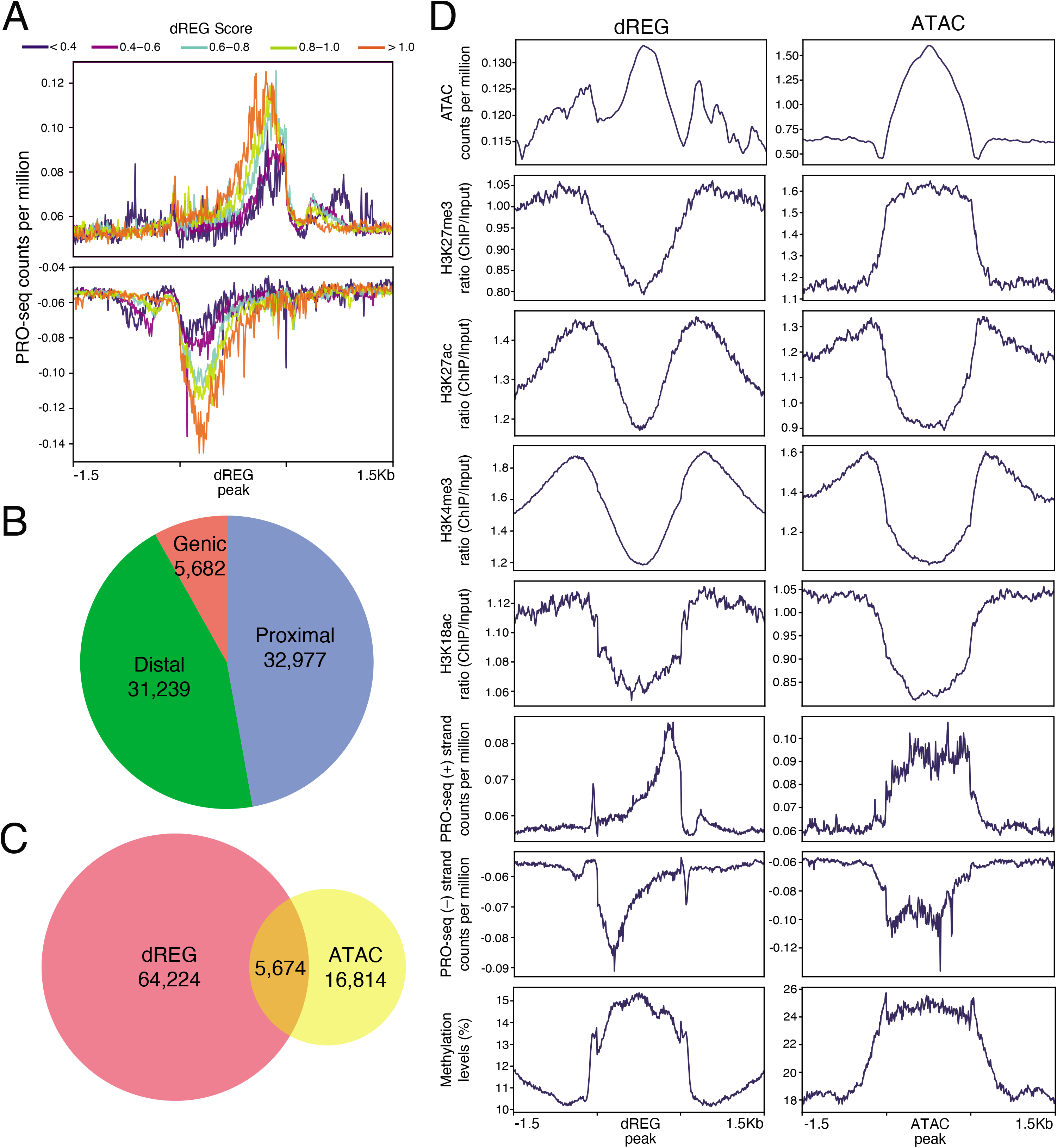
Chromosomal features associated with dREG peaks in the rice genome. (**a**) PRO-seq read counts for positive-sense (top) and negative-sense of dREG peaks. (**b**) Classification of dREG peaks according to the genomic regions it was located. Genic: within coding sequence regions; proximal: within 1 kbp of genic sequences; and distal: more than 1 kbp away from genic sequences. (**c**) Overlap between dREG and ATAC peaks. (**d**) Enrichment of functional genomic sequencing reads 1.5 kbp upstream and downstream of dREG (left) and ATAC (right) peak regions.

The majority of dREG peaks (∼91.9%) were found in intergenic regions (Fig. 1b). We divided the intergenic dREG peaks into two classes depending on their distance to genic sequences: (i) proximal dREG peaks (dREG_proximal_), which are intergenic dREG peaks that are <1 kb to predicted transcription start or end sites, and (ii) distal dREG peaks (dREG_distal_), which are >1 kb away from predicted transcription start or end sites. Within the intergenic regions, we found approximately equal proportions of dREG_proximal_ and dREG_distal_ peaks (Fig. 1b). dREG_proximal_ peaks had significantly higher dREG scores than dREG_distal_ peaks (Mann Whitney U test, p-value <0.0001; Additional file 1: Figure S2).

Many cis-regulatory elements (CREs) that are presumed to be actively engaged have been shown to reside within accessible chromatin regions [26, 55]. We examined the accessibility of dREG peak regions by comparing dREG peaks with open chromatin regions identified using Assays for Transposase-Accessible Chromatin with sequencing (ATAC-seq), and considered a conservative set of peaks that were within 100 bps of each other as overlapping peaks. Of the 69,898 dREG peaks across the genome, we found an overlap of 5,674 (8.1%) dREG peaks with the 23,961 detected accessible chromatin regions (herein referred to as ATAC peaks) as identified by ATAC-seq (Fig. 1c). We then examined any differences in overlap with ATAC peaks for dREG_proximal_ and dREG_distal_ peaks and found that dREG_proximal_ peaks had twice as much overlap with ATAC peaks compared to dREG_distal_ peaks (Additional file 2: Table S1).

In addition to accessible chromatin regions, histone modifications can provide insight into the regulatory mechanisms of the CREs, whether or not they are contained within these accessible chromatin regions. To determine the chromatin features associated with dREG peaks, we used previously generated genome-wide maps of histone modifications and cytosine methylation [19] and compared their profile at ATAC peaks (Fig. 1d). We find that H3K27ac and H3K18ac histone marks were enriched around the dREG and ATAC peaks, (+/- 1.5kb). However, PRO-seq reads showed the expected enrichment of bi-directional transcription activity for dREG peaks but not for ATAC peaks. Moreover, the repressive H3K27me3 mark [56] was enriched within ATAC peaks but depleted within dREG peak regions (Fig. 1d). In addition, the levels of DNA methylation marks [37] were increased for both dREG and ATAC peak sequences, but the latter had higher density of DNA methylation than dREG peaks. We then compared epigenetic marks between dREG_proximal_ and dREG_distal_ peaks and whether they overlapped ATAC peaks or not, and found no overall differences in chromatin features (Fig. S3). Taken together, these results suggest there is a range of detected transcribed regulatory elements (represented by different dREG scores) in the rice genome and that overall they show relatively little overlap with accessible chromatin regions, as they display different signals in terms of their epigenetic architecture. Consistant with our previous work [19], the different combination of epigenetic marks and transcriptional signatures may play a more nunanced role in determining the chromatin state and functionality of a genomic region.

### dREG peaks enriched for DNA methylation overlap with repetitive elements

DNA methylation is often found in inactive regulatory elements, where their target genes are repressed [57], which is why we did not expect to detect an enrichment of DNA methylation for actively transcribed candidate regulatory regions. For instance, when we compared epigenetic marks for transcriptionally active genes with annotated repetitive elements in the rice genome, the common repressive mark H3K27me3 and DNA methylation were enriched within repeat sequences or genes with low expression (Additional file 1: Figure S4). Methylation in plants is dependent on the RNA-directed DNA methylation (RdDM) pathway, wherein transcribed non-coding RNA molecules direct the addition of DNA methylation to specific DNA sequences that are largely associated with transcription repression [58, 59]; thus the PRO-seq data in intergenic regions may, in part, be detecting silencing-related transcription. We therefore investigated whether cytosine methylation was mainly attributed to silencing of repetitive elements or was potentialy an inherent characteristic of plant TREs. We categorized dREG peaks into three repeat classes: (i) “without repeat” class, where the dREG peak region does not contain annotated repeat sequences, (ii) “intermediate repeat” class, where the dREG peak region overlaps with at least 1 bp of a repeat sequence, and (iii) “repeat” class, where the entire dREG peak region overlaps a repeat region. For both dREG_proximal_ and dREG_distal_ peaks, the majority of the peaks (>70%) were in the intermediate repeat class, which had significantly higher dREG scores than both repeat and without repeat class dREG peaks (Mann Whitney U test p-value < 0.001 and Fig. 2a).

**Figure 2.**
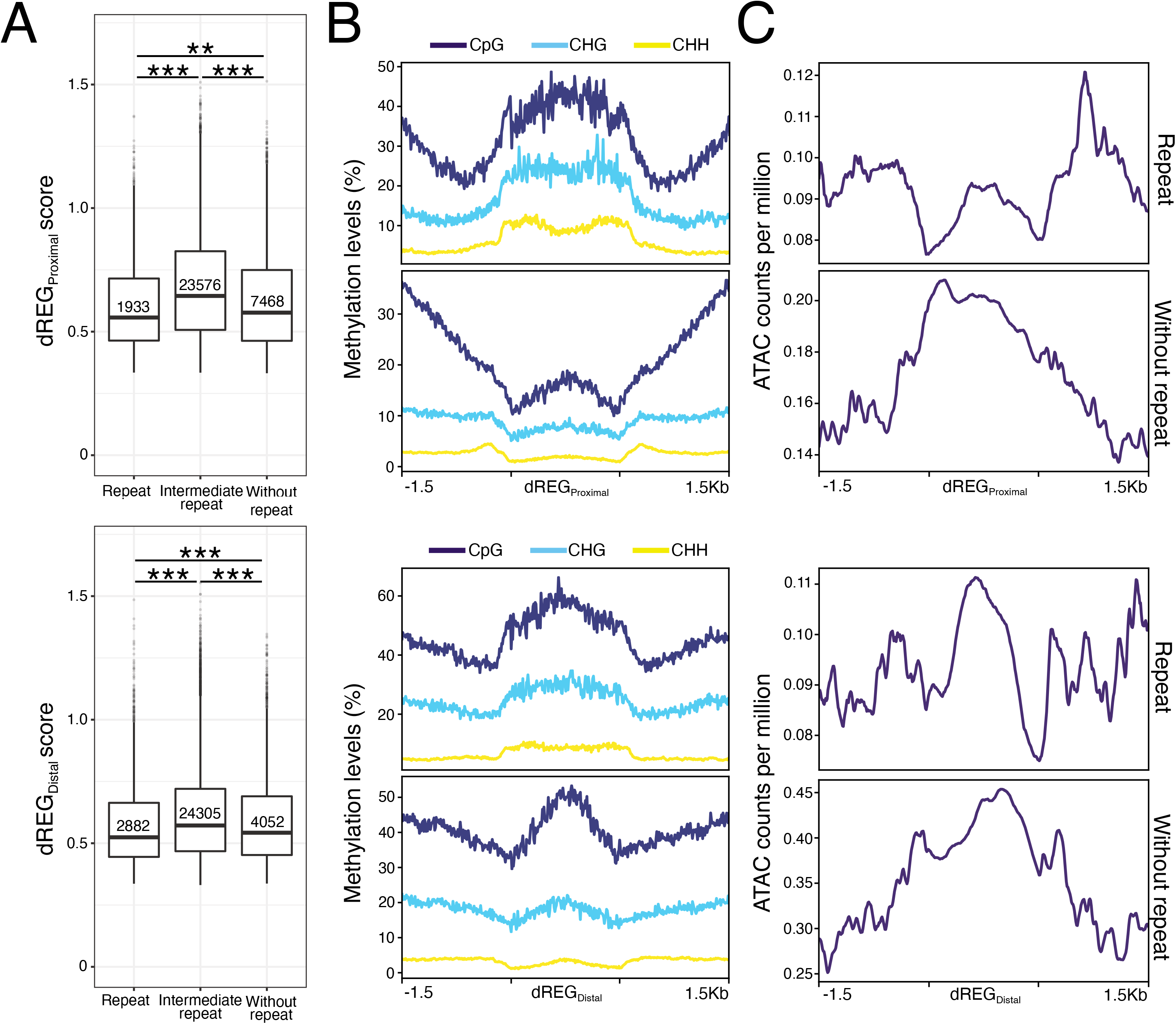
Repetitive sequence characteristics within proximal (top row) and distal (bottom row) dREG peak regions. (**a**) Median dREG scores for dREG peaks classified into three repeat classes: (Left) Repeat: entire dREG peak region is a repetitive sequence; (Middle) Intermediate repeat: dREG peak regions that are not classified as “repeat” or “no repeat” class; (Right) No repeat: no repeat sequence was annotated in dREG peak region. The numbers in the box plot represent the count of dREG peaks in each category. (**b**) Percentage of methylated cytosine for the three different cytosine contexts CpG, CHG, and CHH sites (where H is A, T, or C nucleotide). (**c**) ATAC-seq read counts centered at dREG peak for the repeat classes No repeat and Repeat.

DNA methylation in rice, as in other plants, occurs at three contexts: CpG, CHG, and CHH (where H indicates A, T, or C). We compared these cytosine contexts in dREG_proximal_ and dREG_distal_ peaks, and found that the dREG_distal_ peaks had significantly higher levels of methylation across all three cytosine contexts (Mann Whitney U test p-value < 0.001 and Fig. S5). Also, for both dREG_proximal_ and dREG_distal_ peaks, regardless of repetitive content, CpG sites had the highest methylation level, while CHH sites had the lowest (Fig. 2b). Since a higher dREG score was associated with higher transcriptional activity (Fig. 1a), we plotted dREG peak score and methylation for both dREG_proximal_ and dREG_distal_ peaks. We found that overall methylation levels were negatively correlated with dREG scores (p < 0.01), although CHH methylation, previously associated with TE silencing [36,60–62], had significantly positive correlation (p < 0.001) with dREG scores (Fig. S5).

We then contrasted the methylation profiles of dREG peaks in the repeat and without repeat class, as the former is likely to represent epigenetic marks from chromosomal silencing. When we considered chromatin accessibility, we found that dREG peaks (proximal or distal) in the without repeat class were more accessible (Fig. 2c). Taken together, these results suggest that PRO-seq signals detected by dREG that overlap repetitive elements could be involved in silencing mechanisms (such as RDdM) rather than enhancer activity (candidate active CREs).

### Evidence of selection within dREG peaks

One strategy known for finding candidate functional sequences in genomes is to look for evolutionary constraint across species [43,63–65]. While sequence conservation can identify conserved noncoding sequences (CNSs), other functional CREs, such as enhancers, have been shown to show sequence diversification and be more species specific [66, 67], and therefore not readily detected by conservation-based methods. To infer functionality of our dREG peaks, we estimated the level of evolutionary constraint occurring within the dREG peak regions using two evolutionary-based approaches: phyloP [68] and fitCons [19, 69]. Selection within dREG peaks were compared to selection in coding sequences and neutral regions of the genome (control) (Fig. 3).

**Figure 3.**
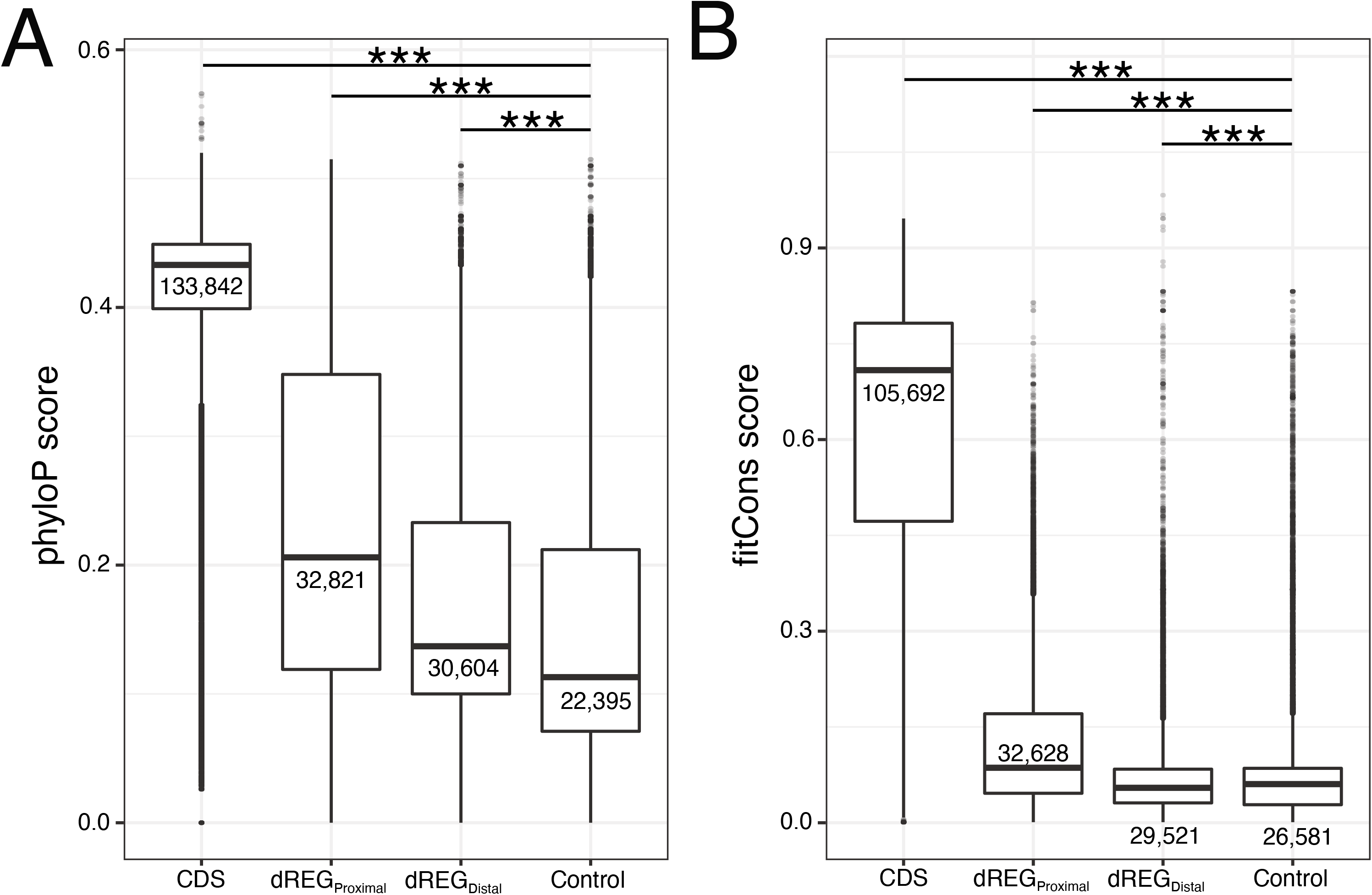
Evolutionary scores. (**a**) phyloP and (**b**) fitcons for dREG peaks and comparison to coding sequence regions or random regions of the genome.

The results showed an overall higher evolutionary constraint at dREG_proximal_ peaks compared to dREG_distal_ peaks. While the peaks had lower levels of purifying selection compared to coding sequences (Mann Whitney U test p < 0.0001), both dREG_proximal_ and dREG_distal_ peaks had higher phyloP values compared to random regions of the genome. The dREG_proximal_ peaks had a significant negative correlation (p-value < 0.001) between dREG score and phyloP statistics, particularly in regions with dREG scores above 1.2 (Additional file 1: Figure S6). A possible explaination for this could be that some of the regions with higher dREG scores are detecting methylation-related activity. For instance, we detect a positive correlation between dREG_proximal_ scores and CHH methylation, which suggests that not all dREG activity is related to transcription but may also be related to silencing activity (Additional file 1: Figure S5).

For fitCons, dREG_proximal_ peaks had higher fitCons scores (ρ) than dREG_distal_ peaks (Mann Whitney U test p < 0.001). For both dREG_proximal_ and dREG_distal_ peaks, we observed a significant positive correlation (p < 0.001) between dREG score and fitCons statistics. We noted that dREG_distal_ peaks had fitCons scores that were significantly lower than dREG_proximal_ and random control region (Fig. 3b). Moreover, lower dREG scores had lower fitCons scores, and lower dREG scores have been attributed to regions evolving more neutrally and within active repetitive elements (Additional file 1: Figure S6) [19, 69].

To investigate why dREG_distal_ peaks had lower fitCons scores, we looked to determine whether we could detect differences in evolutionary constraints at dREG peaks based on their repetitive element content. When we divided dREG peaks by repeat class we found that dREG peaks, regardless of the amount of repeat content, had significantly higher phyloP scores (Mann Whitney U test p < 0.05) than random regions of the genome (Additional file 1: Figure S7). For fitCons scores, compared to random genomic regions, dREG_proximal_ peaks had significantly elevated ρ statistics for all repeat classes (Mann Whitney U test p < 0.001) but for dREG_distal_ peaks, there was no significant difference for the without repeat class, while the intermediate repeat and repeat classes had significantly lower fitCons scores (Mann Whitney U test p-value < 0.0001). A similar trend in dREG peaks was previously observed in rice [19] and human populations [70]; in these cases, there was an excess of sites under weak negative selection was observed, suggesting a recent selection since the most recent common ancestor. Taken together, these results suggest dREG peaks display some level of conservation that is stronger in proximal peaks than distal peaks, although this distinction may be due to the increased occurrence of repetitive elements with the latter.

### Functionality of dREG_proximal_ peaks and gene expression

We explored whether candidate transcribed regulatory elements identified by the dREG algorithm could indeed be functional regulatory elements (active CREs). We first focused on dREG_proximal_ peaks detected in the promoter region and examined whether their activity is associated with expression of nearby genes. The results showed that genes with a dREG_proximal_ peak in its promoter region had significantly higher gene expression (Mann Whitney U test p < 0.001 and Fig. 4a). This higher level of gene expression was observed regardless of the repeat class associated with the peak.

**Figure 4.**
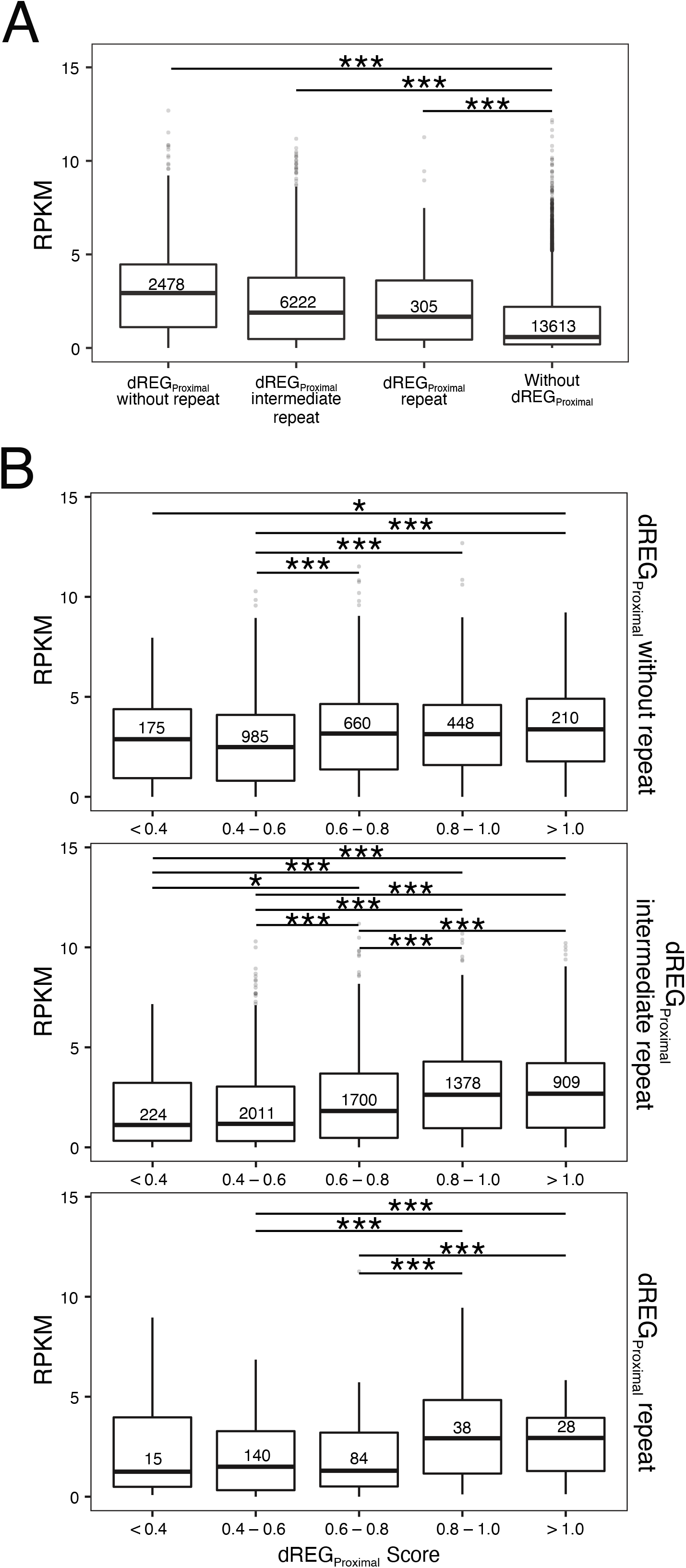
Functional characteristics of dREG_Proximal_ peaks that are detected at 5 prime untranslated regions of genes. (**a**) Expression levels (shown as Reads per kilo base per million mapped reads, RPKM) for genes with and without dREG_Proximal_ peaks. Genes with dREG_Proximal_ peaks were divided by repeat class and their expression levels were compared to the genes without dREG_Proximal_ peaks. Numbers in boxplot represent sample size of genes. (**b**) Gene expression levels for genes with dREG_Proximal_ peaks divided by dREG score and repeat classification. Asterisk (*) indicate significant differences after all pairwise comparisons using the Mann-Whitney U test. Numbers in boxplot represent sample size of genes.

Previous studies using PRO-seq data to define transcribed regulatory elements established a dREG score threshold of >0.8 for human enhancers and < 0.3 for non-functional or random transcriptional activity [54]. In plants, the only dREG threshold that has been explicitely used to characterize these regulatory elements was a dREG score of > 1.0 in the rice genome [19]. Based on this, we examined the relationship between dREG score and functional genomic activity by comparing the relationship between gene expression levels and dREG_proximal_ peak scores. Results showed that genes with a dREG_proximal_ peak and high dREG score (>0.8) had significantly higher gene expression than genes with low (<0.8) dREG score (Mann Whitney U test p < 0.001 and Fig. 4b). This was consistent even in the presence of repeat elements. We also examined the chromatin profiles surrounding dREG_proximal_ peaks and discovered that peaks with scores higher than 0.8 had similar chromatin marks (Additional file 1: Figure S8), although with some unexpected increase in DNA methylation levels in all cytosine contexts. The methylation pattern in peaks with higher dREG scores appears different from those with lower scores. dREG peaks with higher scores display methylation in the center of the peak, in contrast to methylation being distributed throughput the lower dREG scored peaks. Methylation was therefore taken into consideration in our subsequent downstream analysis.

### Identifying distal transcribed regulatory elements through bidirectional transcription activity

Distal transcribed regulatory elements associated with dREG_distal_ peaks may signal RNA-producing enhancers, producing unstable transcripts in both directions [71]. Indeed, in animals, genomic coordinates of distal transcribed regulatory elements have often overlapped with enhancers that actively produce eRNAs, although in plants there is no evidence that these are indeed associated with enhancer activity.

To further characterize rice dREG_distal_ peaks and identify potential functional roles in the rice genome, we examined long-range chromosomal contact interaction between genes and candidate CREs using the Pore-C sequencing method [72]. Pore-C couples chromatin conformation capture with long-read nanopore sequencing to detect genome-wide multi-way chromatin contacts. It has been shown to be highly effective at sequencing through repeat regions of the genome and is able to detect increased contact intensity with less sequencing reads than conventional Hi-C sequencing [72]. Using Pore-C sequencing on rice leaves, we generated 104 million concatamer reads that correspond to 290 million contacts across the rice genome (Additional file 2: Table S2).

Since Pore-C sequencing has not been applied in plants to profile chromatin activity, we first examined the functionality of the 3D genome architecture detected by this method. Using the Pore-C method, we detected 3,261 distinct topologically associated domains (TADs) which are localized chromosomal regions of high physical contact [73]. TADs had a median size of 80 kb and the TAD boundaries were enriched for transcription and active chromatin marks (Fig. 5a and Fig.S9). These results were consistent with previous *Oryza* Hi-C sequencing results [74] indicating that Pore-C sequencing was able to detect functional 3D contacts across the *Oryza* chromosome.

**Figure 5.**
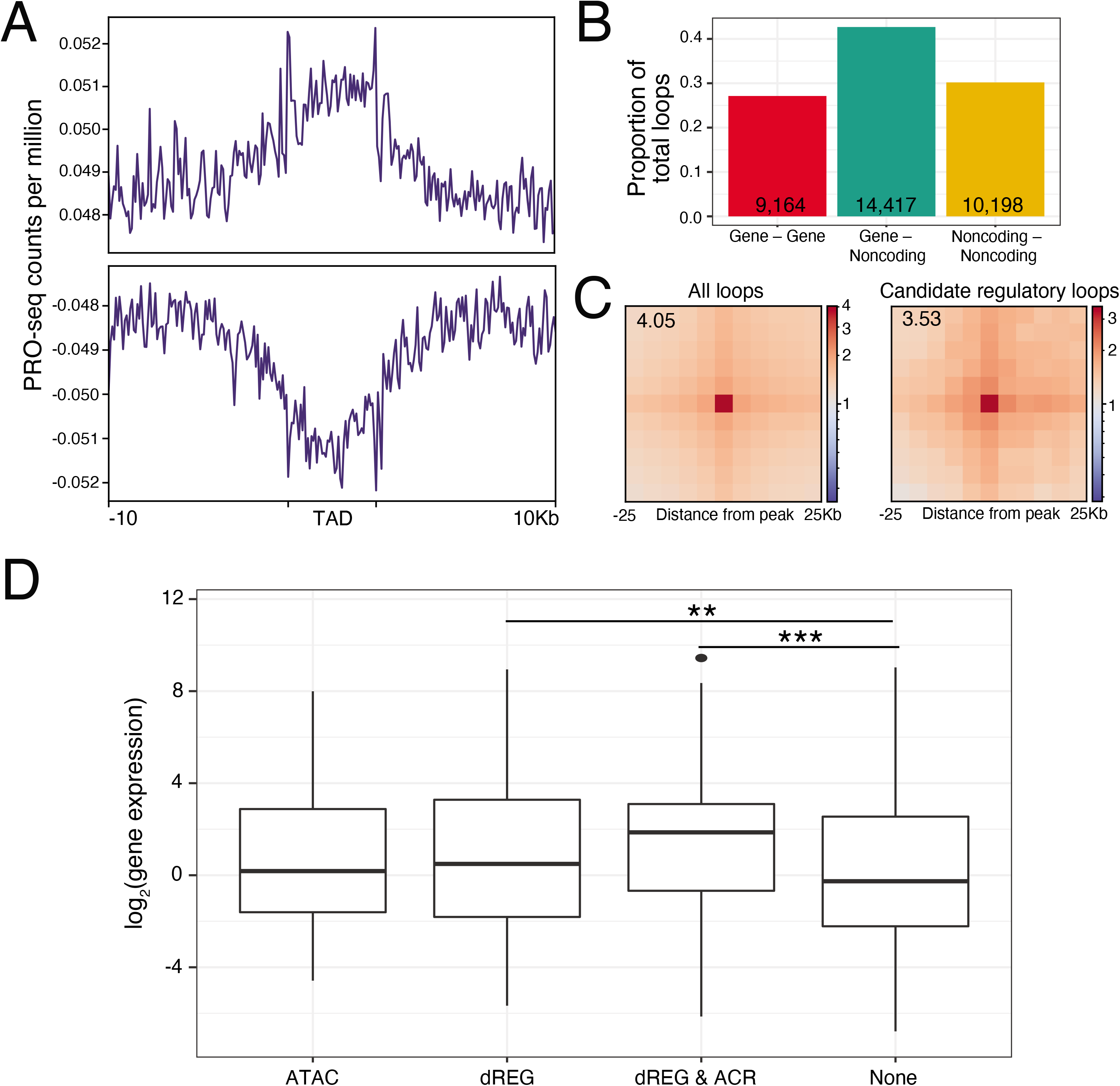
Chromatin features and functionality of dREG_Distal_ peaks. (**a**) PRO-seq read count enrichment surrounding TADs. Shown are 10 kbp upstream and downstream of TADs with the TAD scaled to 5 kbp. PRO-seq read counts were averaged in 100 bp windows. (**b**) Proportion of gene-gene, gene-noncoding, and noncoding-noncoding loops that were detected using Pore-C sequencing. (**c**) Aggregate Peak Analysis (APA) plots showing the aggregated Pore-C contacts around chromatin loops identified in all chromosomes (left) and only between a noncoding-gene loop (right). The plot is a pile-up of 25 kbp upstream and downstream of loop anchors (centered in each axis) of every identified loop. Color represents log2 fold enrichment of the observed aggregated matrix over a normalization matrix that was aggregated from randomly shifted controls regions across the chromosome. (**d**) Gene expression level for genes that are contacting a noncoding anchor with either an ATAC peak, dREG peak, or has no annotation. Asterisk (**) indicates a p-value < 0.01 and *** indicates a p-value < 001 after an FDR-corrected Mann-Whitney U test.

To determine whether dREG_distal_ peaks could represent distal regulatory elements of one or multiple target genes, we used the Pore-C sequencing data to detect the formation of chromatin loops at 5 kbp resolution. Using the FitHiC2 algorithm [75], we detected 33,779 chromatin loops, where 42.7% (14,417) of those loops involved a gene (where the coding window had to contain more than 100 bp of coding sequences) and a noncoding region (Fig. 5b). These represent candidate gene-regulatory element loop interactions. We then visualized this loop interaction by conducting aggregate peak analysis (APA), which takes the contact map and measures its enrichment with respect to its local neighborhood (signal around detected chromatin loop formations) (Fig. 5c). When we compared the APA plots for all chromatin loops and candidate gene-regulatory element loops, the latter loops had strong enrichment of signals centered at the contact point. Specifically for the candidate gene-regulatory element loops, the central window (*i.e.* the contact point) had ∼3.5 fold increased in contacts compared to the lower left windows (*i.e.* background contact levels) (Fig. 5c).

Next, we examined the potential functionality of candidate gene-regulatory element loops. As a first step, we processed a subset of candidate loops as probable gene-regulatory element interactions (see the “Methods” section for details). To do so, we (i) focused on chromatin loops that do not cross TAD boundaries, as transcription related chromatin loops co-localize within TAD domains in plants [76], (ii) removed loops where the noncoding anchor had peaks with CHH methylation, as these are more likely to represent transposable element silencing activity [37], and (iii) removed loops where the gene anchor overlapped multiple genes, as the resolution of our Pore-C data could not differentiate whether the noncoding region regulated one or multiple genes.

Following these filtering steps, we analyzed gene-regulatory element loops where the noncoding anchor had only an ATAC peak, only a dREG peak, or had both an ATAC and dREG peak (Fig. 1c). These loops were then compared to loops where the noncoding anchor had no detected ATAC peak nor dREG peak and represented control gene-noncoding sequence interactions (Additional file 1: Figure S10A). We found that for all post-filtered gene-noncoding sequence loops, and regardless of the annotation within the noncoding anchor, the majority of genes had contact with a single noncoding region (Additional file 1: Figure S10B). Furthermore, the average distance between the gene and the noncoding region was 65 kb to the dREG only peak, 45 kb to the ATAC only peak, 35 kb to both dREG and ATAC peaks, and 50 kb to a noncoding anchor with no annotation (Additional file 1: Figure S10B). We also found that genes that contacted a dREG peak had significantly higher gene expression than those that contacted a noncoding anchor with no annotation (Mann Whitney U test p < 0.01 and Fig. 5d). In contrast, genes that contacted an ATAC peak did not show any significant difference in expression. Finally, genes with both dREG and ATAC peaks had significantly elevated gene expression (Mann Whitney U test p-value < 0.001 and Fig. 5d). The median dREG scores for gene-contacting dREG peaks were 0.54 (Additional file 1: Figure S11). Gene ontology enrichment analysis on genes found in gene-non coding loops did not highlight specific pathways or function.

### dREG_distal_ peaks are targets of transcriptional regulation

As an orthogonal approach to investigate the gene regulatory functions of dREG peaks, we identified expression quantitative trait loi (eQTLs) across a panel of rice varieties and intersected those eQTLs with dREG peak regions. Using gene expression and SNP data from 216 rice varieties grown in well-watered field conditions [77] we detected 274,480 eQTLs after a 5% Bonferroni threshold. We intersected the dREG_distal_ peaks with the significant eQTLs and found an overlap with 13,036 eQTLs.

To test the significance of the observed overlap, we generated a bootstrap distribution by randomly sampling the potentially non-functional regions of the genome, which were matched for size and total number of dREG_distal_ peaks. Results showed that the observed number of overlap between eQTLs and dREG_distal_ peaks (Fig. 6) was higher than the maximum number of overlaps in the bootstrap distribution across random regions, indicating dREG_distal_ peaks are enriched for eQTLs. We also examined dREG_distal_ peaks that were limited to those forming loops with genes and potentially involved in transcription (identified and analyzed in Fig 5d). Overlap of those filtered dREG_distal_ peaks were also significantly enriched for eQTLs (Additional file 1: Figure S12). We repeated the process for distal ATAC peaks (ATAC_distal_) that are intergenic and more than >1 kb away from predicted transcription start or end sites, and found a significant under representation for eQTLs (Additional file 1: Figure S13). Taken together, these results suggest that dREG_distal_ peaks could be good candidate TREs that could impact gene activity by promoting transcription.

**Figure 6.**
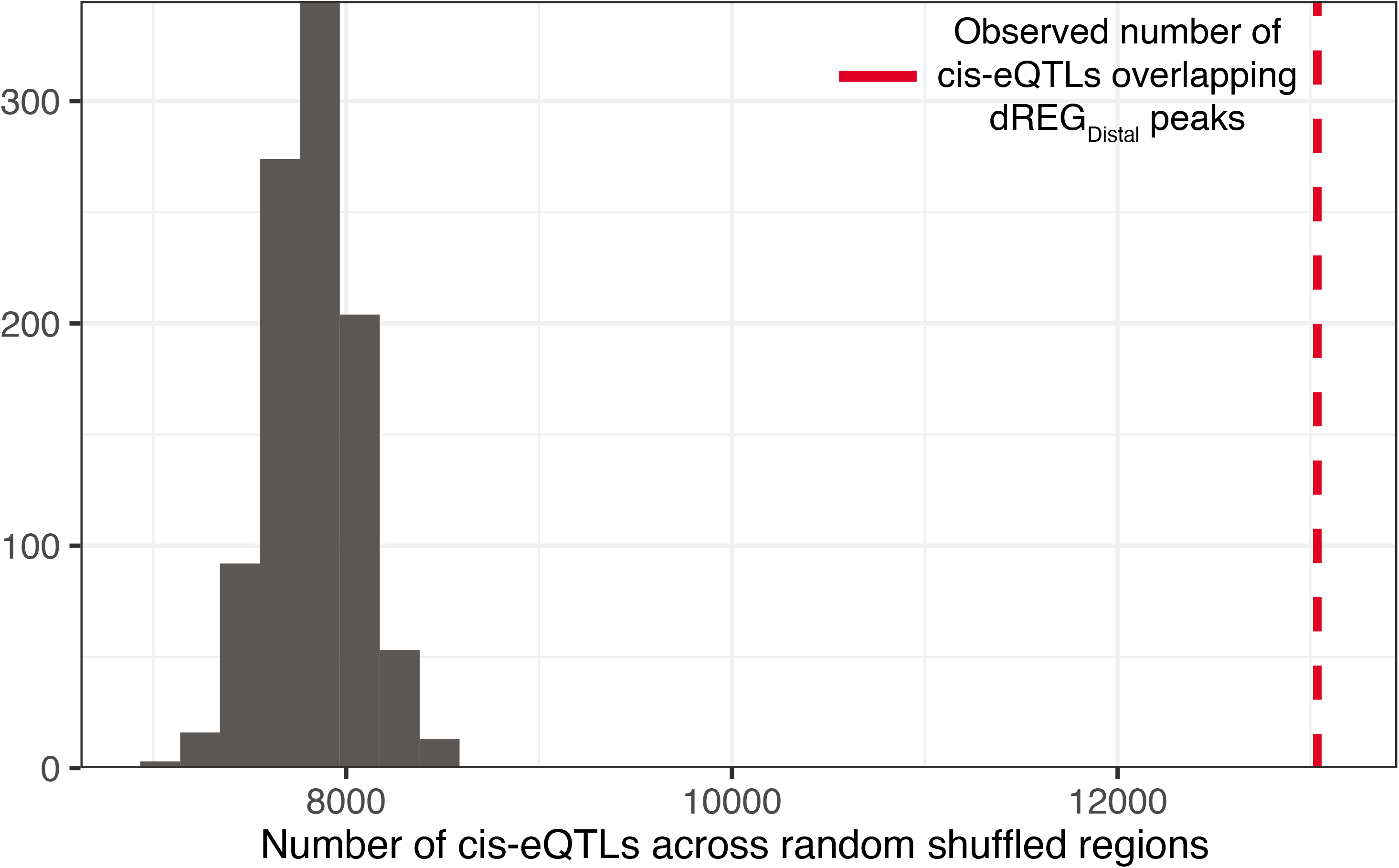
Enrichment of eQTLs within dREG_distal_ peaks. Histogram shows the bootstrap distribution of the total number of eQTLs overlapping a random region of the genome that are matched for size and total number of dREG_distal_ peaks. The red dotted line shows the total number of eQTLs overlapping the dREG_distal_ peaks.

### Summary: nascent transcription and the rice genome

The rice genome encodes a large number of transcripts that require spatio-temporal regulation and for which a large of *cis*-regulatory elements remain to be characterized. In this study, we took advantage of complementary functional genomics datasets to characterize with high sensitivity the synthesis of eRNAs that arise at both distal and proximal genomic elements. We considered a broad range of dREG scores (from 0.3 to >1.0) to explore a wider range of transcribed regulatory elements that could have distinct functional roles based on their functional genomic signatures. We find that dREG peaks (proximal and distal) that did not overlap with repetitive elements exhibited greater evolutionarily constraint and had a higher incidence of overlap with ATAC peaks. In addition, dREG_proximal_ peaks with higher dREG scores correlated with increased transcription of nearby genes, and gene expression was higher for proximal peaks that did not overlap with repeat sequences (Figs. 2 and 4).

Studies in plant genomes, and particularly in maize, have relied mainly on accessible chromatin regions (e.g. ATAC peaks) as an indicator of the presence of active distal CREs [26, 74]. In study by Lozano et al. [18] using dREG/PRO-seq data in maize, they found that 31% of their identified TREs co-located with a list of distal ATAC peaks characterized by Ricci et al. [26], as well as an overlap of 17% between their list of intergenic regulatory elements and CREs found to form CRE-gene loops. Our results suggests a much lower co-localization between dREG and ATAC peaks, with an overlap of about 3.7%. This could be due in part to differences between species in their 3D chromatin architecture, as has been reported with the growing number of chromosomal contact experiments in plants [78]. Our results suggest that a proportion of dREG_distal_ peaks could be functional distal transcribed regulatory elements, as we detected a significant increase in gene expression in gene-noncoding loops that contain a dREG peak compared to gene-noncoding loops with no annotation or even with only an ATAC peak. However, since the loops with both an ATAC and a dREG peak is associated with the highest gene expression levels, we still have to determine which elements are functional, whether it requires the overlap between eRNA (i.e., bidirectional transcription) activity and chromatin accessibility, or whether the presence of eRNAs is sufficient.

One of the key observations in this study is the impact of the presence of repetitive elements overlapping the detected dREG peaks. For dREG peaks that partially or entirely overlapped with annotated repetitive elements, we detected a higher percentage of methylation across all three cytosine contexts. Interestingly, we observed an increase in CHH methylation, which occurs predominantly at TEs, and has been shown to be involved in the prevention of transposon jumping during development in Arabidopsis [79]; this methylation occurs through the plant-specific RdDM pathway that operates via non-coding RNA [80]. The fact that DNA methylation can positively or negatively impact transcriptional activity, for example by modulating binding affinity of TFs, makes it a confounding factor when characterizing candidate TREs. In plants, we argue that methylation profiles should always be considered with PRO-seq datasets to characterize the intergenic transcription signal. We also note that the dREG method uses a training set of mammalian transcribed regulatory elements, which may not be optimal for plant genomes. We cannot rule out that there may have been a level of RdDM transcription contamination in the set of transcribed regulatory elements, but this was taken into account when we filtered out dREG sites presenting methylation.

Beyond the potential presence of TE silencing mechanism associated with dREG peaks, we noted that dREG scores were significantly higher for the intermediate repeat class, but this class may contain different types of elements, as an overlap of as little as 1 bp with a repetitive element places a dREG peak in this category. An interesting perspective could be the presence of an overlap between dREG peaks and *cis*-regulatory elements derived from transposable elements, as a result of an evolutionary process called TE exaptation, or TE co-option [81–83]. In mammals, there are several reported instances of TEs providing CREs, including enhancers and repressive elements, and TEs have contributed an important fraction of TF binding sites across the genome (5–40% [84, 85]). While in plants the contribution of TEs for CREs is less clear, future characterization of these overlapping regions could be an interesting avenue to identify potential cases of TE-derived CREs. Overall, this suggests that considerations, such as methylation levels and the potential differences with chromatin accessibility, have to be taken into account when addressing transcribed regulatory elements in plants.

## Conclusions

In conclusion, we have characterized eRNA producing regions in the rice genome. We find that some of these share features of enhancers and are associated with transcription regulation, which makes them compelling candidate enhancer elements. While the production of eRNAs may be considered a key characteristic used for identifying enhancers in animal studies, there remains a debate as to whether every enhancer (in animal systems) produces eRNAs, even at low levels that are not detected by current methods [51]. In this study, we used the assumptions that eRNAs act principally in *cis* versus *trans*, due to the relative instability of eRNAs [8, 67] and several studies have demonstrated eRNA-dependent transcriptional regulation of mRNAs produced from loci adjacent to the corresponding eRNA-producing regions [12,86,87]. Further characterization of eRNA producing regions in other plant genomes will help us better understand whether this assumption holds true for plants.

## Materials and Methods

### Plant material

Seeds of *O. sativa* landrace Azucena (IRGC 328; tropical Japonica), provided by the International Rice Research Institute (Los Baños, Philippines), were used for the functional genomic datasets. Seeds were incubated for 5 d at 50° C and germinated in water in the dark for 48 h at 30° C. These were subsequently sown on hydroponic pots suspended in 1× Peters solution and 1.8 mM FeSO4 (pH = 5.1–5.8) (JR Peters). Plants were grown for 15 d in growth chambers (12-h days; 30 °C/20 °C day/night; 300–500 μmol quanta m−2 s−1; relative humidity: 50–70%). Leaf tissue for library construction was collected from 17-d-old, young plants.

### RNA-Seq

Total RNA was extracted using RNeasy Plant Mini kits (Qiagen), according to the manufacturer’s instructions. RNA quality was determined by BioAnalyzer (Agilent). Contaminating DNA was removed from total RNA samples with Baseline-ZERO DNase (Epicentre), whereas ribosomal RNA was removed using a Ribo-Zero rRNA Depletion Kit (Epicentre). Strand-specific RNA-Seq libraries were synthesized using a Plant Leaf ScriptSeq Complete Kit (Epicentre). Libraries were sequenced for 2 × 100-bp reads on an Illumina HiSeq 2500. Two biological replicates were generated and a third replicate (SRA : SRX7082160; Bioproject : PRJNA586887) generated under the same conditions and used in a previous study [19] was used. The sequencing reads were adapter-trimmed and quality-controlled using BBTools (https://jgi.doe.gov/data-and-tools/bbtools/) bbduk program version 37.66 with option: minlen = 25 qtrim = rl trimq = 10 ktrim = r k = 25 mink = 11 hdist = 1 tpe tbo. Trimmed reads were aligned to the Azucena reference genome [45] (Bioproject PRJNA424001) using hisat2 version 2.2.1 [88] and estimated the read counts for each gene using featureCounts [89]. To normalize the variation existing between different samples, we applied the trimmed mean of M value (TMM) method [90] from the edgeR version 3.18.0 package [91] on each samples’ gene expression values. For each gene the expression values were averaged across the three replicates.

### DNA methylation

DNA was extracted using DNeasy Plant Mini kits (Qiagen) following the manufacturer’s protocol. Extracted DNA was sheared into 350-bp fragments using an S220 Focused-ultrasonicator (Covaris). An Illumina TruSeq DNA Kit (Cat. No. FC-121-3001) was used to construct the library and a Zymo Lightning Kit (Cat. No. D5030) was used to perform the bisulfite treatment. KAPA Uracil Polymerase (Cat. No. KK2623) was used to amplify the library with 12 cycles. One biological replicate was generated and a second replicate, generated under the same conditions and used in a previous study [19] was used (SRA : SRX7082155; Bioproject : PRJNA586887). Libraries were sequenced using Illumina protocols for 2×100-bp reads on an Illumina HiSeq 2500. Raw bisulfite sequencing (BS-seq) reads quality controled using the program trim galore Ver. 0.6.6 (http://www.bioinformatics.babraham.ac.uk/projects/trim_galore/) with default parameters. We used bismark version 0.16.3 [92] for mapping the BS-seq reads and deduplicating reads.

### Chromatin accessibility

Intact nuclei were isolated using the plant nuclei isolation protocol described by Zhang and Jiang [38]. Nuclei quality was assessed using DAPI staining. Chromatin was fragmented and tagged following the standard ATAC-seq protocol [93]. Libraries were purified using Qiagen MinElute columns before sequencing and were sequenced as paired-end 51-bp reads on an Illumina HiSeq 2500 instrument. Sequencing reads were adapter trimmed and QC controlled using the script bbduk.sh version 38.90 (https://sourceforge.net/projects/bbmap/) with parameters: minlen=16 qtrim=rl trimq=20 ktrim=r k=19 mink=10 hdist=1 tpe tbo. Trimmed sequencing reads were aligned to the Azucena reference genome using Bowtie 2 version 2.4.2 [94] under option very-sensitive and with the parameter -X 1000. The Azucena reference genome included the chloroplast sequence (genbank ID: GU592207.1) to allow chloroplast originating ATAC-seq reads, and were subsequently removed using the script removeChrom.py from the Havard FAS informatics group (https://github.com/jsh58/harvard/blob/master/removeChrom.py). Peak calling were conducted using the program MACS2 version 2.2.7.1 [95] with the parameters: --nomodel -g 379627553 -f BED -q 0.05 --extsize 200 --shift -100 --keep-dup all -B. We used MACS2 to call peaks for each of the three replicate libraries and peaks that overlapped 50% in size between at least two replicates were chosen for downstream analysis. To determine ATAC peaks that overlapped dREG peaks we used bedtools closest function and peaks that were within 100 bp of each other were considered as overlapping peaks.

### ChIP-Seq

Leaf tissue (2 g) was fixed in 1% formaldehyde (v/v) for 15 min, after which glycine was added to a final concentration of 125 mM (5 min incubation). Tissues were rinsed three times with de-ionized water before being flash frozen in liquid nitrogen. Chromatin extraction and chromatin shearing were performed using a Universal Plant ChIP-seq kit (Diagenode) following the manufacturer’s instructions. Protease inhibitor cocktail (MilliporeSigma) was added to extraction buffer. Samples were sonicated for 4 min on a 30 s on/30 s off cycle using a Bioruptor Pico (Diagenode). Subsequent steps were performed as in the Universal Plant ChIP-seq kit protocol. Immunoprecipitation was done using anti-acetyl-histone H3 (Lys27) (H3K27ac; Cell Signaling Technology; Cat. No. 4353S; lot 1), anti-trimethyl-histone H3 (Lys27) (H3K27me3; MilliporeSigma; Cat. No. 07-449; lot 2919706), anti-trimethyl-histone H3 (Lys4) (H3K4me3; EMD Millipore; Cat. No. 07-473; lot 2746331) and anti-acetyl-histone H3 (Lys18) (H3K18ac; Cell Signaling Technology; Cat. No. 9675S; lot 1). The quality and fragment size of immunoprecipitated DNA and input samples were measured using agarose gel electrophoresis and TapeStation 2200 (Agilent). Libraries were synthesized using a MicroPlex Library Preparation Kit (v.2; Diagenode). Libraries were sequenced as 2 × 50-bp reads on an Illumina HiSeq 2500 instrument. Two biological replicates were generated and a third replicate, generated under the same conditions and used in a previous study [19] was used (SRA : SRX7082158 H3K4me3; SRX7082157 H3K18Ac; SRX7082156 H3K27Ac; SRX7082153 H3K27me3; Bioproject : PRJNA586887).

Sequencing reads were adapter trimmed and QC controlled using the script bbduk.sh ver. 38.90 (https://sourceforge.net/projects/bbmap/) with parameters: minlen=16 qtrim=rl trimq=20 ktrim=r k=19 mink=10 hdist=1 tpe tbo. Trimmed reads were aligned to the Azucena reference genome [45] (Bioproject PRJNA424001) using Bowtie 2 version 2.4.2. [94] under option very-sensitive and with the parameter -X 1000.

### PRO-Seq

Nuclei isolation was as described by Hetzel et al. [20], with some modifications. ∼20 g of leaf tissue from 17-d-old plants was collected in 4 °C, placed in ice-cold grinding buffer and homogenized using a Qiagen TissueRuptor. Samples were filtered and pellets were washed twice, followed by homogenization, resuspension in storage buffer (10 mM Tris (pH 8.0), 5 mM MgCl2, 0.1 mM EDTA, 25% (v/v) glycerol and 5 mM DTT) and freezing in liquid nitrogen. Nuclei were stained with DAPI and loaded into a flow cytometer (Becton Dickinson FACSAria II). Around 15 million nuclei were sorted based on the size and strength of the DAPI signal, and subsequently collected in storage buffer. Nuclei were pelleted by centrifugation at 5,000g and 4 °C for 10 min, and resuspended in 100 μl storage buffer.

PRO-Seq was performed as described by Mahat et al. [16], generating strand-specific libraries with reads starting from the 3′ end of the RNA. Amplified libraries were assessed for quality on a TapeStation before sequencing with 1 × 50-bp reads on a HiSeq 2500. One biological replicate was generated and a second replicate, generated under the same conditions and used in a previous study [19] was used (SRA : SRX7082159; Bioproject : PRJNA586887).

The raw reads were then used on the proseq2.0 (https://github.com/Danko-Lab/proseq2.0) pipeline [50] that automatically pre-processes the reads, aligns to the reference genome, and generates output bigWig files for downstream PRO-seq peak calling analysis. To identify peaks of divergent transcription activity we used the bigwig file generated from the previous step as an input for the cloud computing version of the dREG algorithm (https://dreg.dnasequence.org/).

### PoreC data generation and computational processing

We generated PoreC libraries following the protocol of Choi et al. [96] and sequencing library was prepared using the Oxford Nanopore Technologies standard ligation sequencing kit SQK-LSK109. Sequencing was conducted on a GridION X5 and PromethION sequencer and the raw data were base-called by Oxford Nanopore Technologies basecaller Guppy version 4.4.0 (available on https://community.nanoporetech.com/) on high-accuracy mode. The Pore-C data analysis was conducted using the PoreC snakemake workflow developed by Oxford Nanopore Technologies (https://github.com/nanoporetech/Pore-C-Snakemake). Briefly, the pipeline first aligns the nanopore Pore-C chromosome contact sequence reads to the Azucena genome using bwa-sw version 0.7.17-r1188 [97] with parameters -b 5 -q 2 -r 1 -T 15 -z 10 The alignment BAM file was processed with Pore-C tools (https://github.com/nanoporetech/pore-c) to filter spurious alignments, detect ligation junctions, and assign fragments that originated from the same chromosomal contacts. The workflow also generates cool and hic files that can be used for downstream analysis.

### TAD and chromatin loop calling

The PoreC contact matrix generated from the previous analysis was normalized using the KR algorthim [98] with the computational suite HiCExplorer version 3.4 [99]. Using the normalized contact matrix the algorithm topdom [100] was used to call TADs as the method was shown recently to be a highly effiecnt and accurate method for detecting TADs [101]. The topdom analysis was conducted using a 5 kbp resolution contact matrix. PoreC contact matrix was also used to statistically determine the significant chromatin contacts using the program FitHiC2 [75]. Using the genomic distance between windows and their contact probability, FitHiC2 applies a spline fit to model an empirical null distribution and detect chromatin contacts as outliers to this null distribution. FitHiC2 was run with default parameters using the 5kbp resolution contact matrix, while setting the lower bound on the intra-chromosomal distance range (parameter -L) as 10 kbp and upper bound (parameter -U) as 1 Mbp. Candidate chromosome loops were filtered by selecting for window pairs that had a Benjamini-Hochberg procedure based false discovery rate threshold q-value < 0.05. Window pairs that had significant evidence of contact were then classified as whether it was a coding or noncoding window by defining a coding window as those that contained more than 100 bp (i.e. greater than 2% of the window) of coding sequences.

### Azucena reference genome repeat sequence and gene annotation

Repetitive sequences in the Azucena genome were annotated using the EDTA program [102]. The Azucena reference genome lacked gene annotation. To annotate the gene we took the gene models from the Nipponbare reference genome, which arguably has the best gene models for rice, and lifted over the gene coordinates using the program liftoff [103].

### Evolutionary analysis

To calculate phyloP scores we first generated whole genome alignments of wild rice (*O. nivara, O. rufipogon, O. punctata, O. glaberrima, O. barthii, O. brachyantha, O. glumaepatula, O. meridionalis*, and *Leersia perrieri*) [104]. The wild rice reference genomes were aligned to the repeat masked Azucena reference genome using LASTZ version 1.03.73 [105]. Alignment blocks were chained and filtered using the UCSC Kent utilities suite (http://hgdownload.cse.ucsc.edu/admin/exe/linux.x86_64.v287) to obtain a single chain the highest score to represent a single orthologous region of the reference genome. A final multi-genome alignment was generated using the aligner MULTIZ [106]. Using the multi-genome alignment four-fold degenerate sites were extracted using the phast version 1.3 package [107]. The four-fold degenerate sites were then used to build a phylogenetic tree using raxml version 8.2.12 [108] with the GTR gamma model. The topology obtained from the phylogenetic analysis and the four-fold degenerate site data was used to fit a phylogenetic neutral model with phylofit. Using the neutral model we estimated the per-base conservation score using the phylop program with mode CONACC and method LRT.

The fitCons score were obtained from Joly-Lopez et al. [19]. But because the fitCons score were calculated using the Nipponbare reference genome we converted those scores to Azucena reference genome coordinates, by aligning the Azucena reference genome to Nipponbare reference genome and using the program liftOver from the Kent utilities suite.

### eQTL detection

Population whole genome resequencing and gene expression data were obtained from Groen et al. [77]. For genes that had multiple transcript expression profile, we chose the longest transcript to represent the expression level of that gene. We conducted eQTL analysis using the program MatrixeQTL [109]. To account for population structure we used plink [110] to calculate structure using polymorphism data and chose the first 5 principal components as covariates to the eQTL model. Resulting p-values for each SNP were filtered using Bonferroni correction and SNPs with adjusted p-value < 0.05 were considered significant eQTLs.

### Gene ontology and motif enrichment

Gene ontology (GO) analysis of genes in gene-non coding loops was performed using BinGO [111] with the full list of GO terms (GO_Full) or using PANTHER [112] with the molecular functions, biological process and cellular component GO lists. Motif enrichment was determined using Homer (v.4.10; http://homer.ucsd.edu/homer/) findMotifs with the options -mset plants -len 6,7,8 enabled, and permuted sets of input sequences were used as controls.

### Plotting functional genomic data

Enrichment of functional genomic reads around peaks of interest were plotted using deeptools [113], specifically the program computeMatrix. APA plots were generated using the program coolpup.py [114].

## Declarations

### Ethics approval and consent to participate

Not applicable.

### Consent for publication

Not applicable.

### Availability of data and materials

All raw sequencing data generated from this study are uploaded on the NCBI Sequence Read Archive. The functional genomic data RNA-seq, ChIP-seq, BS-seq, and PRO-seq that were generated in this study was uploaded under PRJNA586887 specifically with the identifiers XXXXXX-YYYYYYY. For PoreC the FAST5 files are available under accession numbers SRR13985185, SRR13985186, SRR13985187, SRR13985192, SRR13985193 and the FASTQ files are available under accession numbers SRR13985180, SRR13985181, SRR13985182, SRR13985190, SRR13985191.

### Contributions

JYC, ZJL, and MDP designed the idea and participated in revision and discussion; ZJL, JYC, and AEP did the data collection; JYC and ZJL executed data analysis; AJ helped in data analysis; ZJL and JYC drafted the manuscript with significant participation of MDP. All author(s) read and approved the final manuscript.

## Acknowledgements

We would like to thank David Xiaoguang Dai, Priyesh Rughani, Scott Hickey, Eoghan Harrington, and Sissel Juul from Oxford Nanopore Technologies for their assistance on the PoreC data generation. We thank the New York University Center for Genomics and Systems Biology GenCore Facility for sequencing support.

## Corresponding author

Correspondence to Michael D. Puruggangn or Zoé Joly-Lopez.

## Additional files

### Additional file 1: Supplementary figures

**Supplemental Fig 1.** Distribution of genome-wide dREG scores.

**Supplemental Fig 2.** dREG scores for dREG_proximal_ and dREG_distal_ peaks.

**Supplemental Fig 3.** Epigenetic marks for dREG_proximal_ and dREG_distal_ peaks.

**Supplemental Fig 4.** Epigenetic marks for coding and repetitive sequences in the rice genome.

**Supplemental Fig 5.** DNA methylation levels for dREG_proximal_ and dREG_distal_ peaks.

**Supplemental Fig 6.** Scatter plot for dREG peak regions’ score and evolutionary conservation scores.

**Supplemental Fig 7.** Evolutionary scores for dREG_proximal_ and dREG_distal_ peaks.

**Supplemental Fig 8**. Chromatin profiles of dREG_Proximal_ peaks that are binned by dREG scores.

**Supplemental Fig 9.** Epigenetic marks surrounding TAD boundaries.

**Supplemental Fig 10.** Distribution of loops detected by Pore-C for dREG_distal_ peaks.

**Supplemental Fig 11.** Distribution of dREG scores for the dREG peaks contacting a gene.

**Supplemental Fig 12.** Enrichment of eQTLs within dREG_distal_ peaks identified in Figure 5D.

**Supplemental Fig 13.** Enrichment of eQTLs within ATAC peaks.

### Additional file 2: Supplementary tables

**Supplemental Table 1.** Total number and proportion of genic, distal, and proximal dREG peaks that overlapped a ATAC peak region.

**Supplemental Table 2.** Pore-C summary statistics.

