## Supplemental Figures for "Nascent transcription and the associated *cis*-regulatory landscape in rice"

This file includes:

Figures S1 to S13

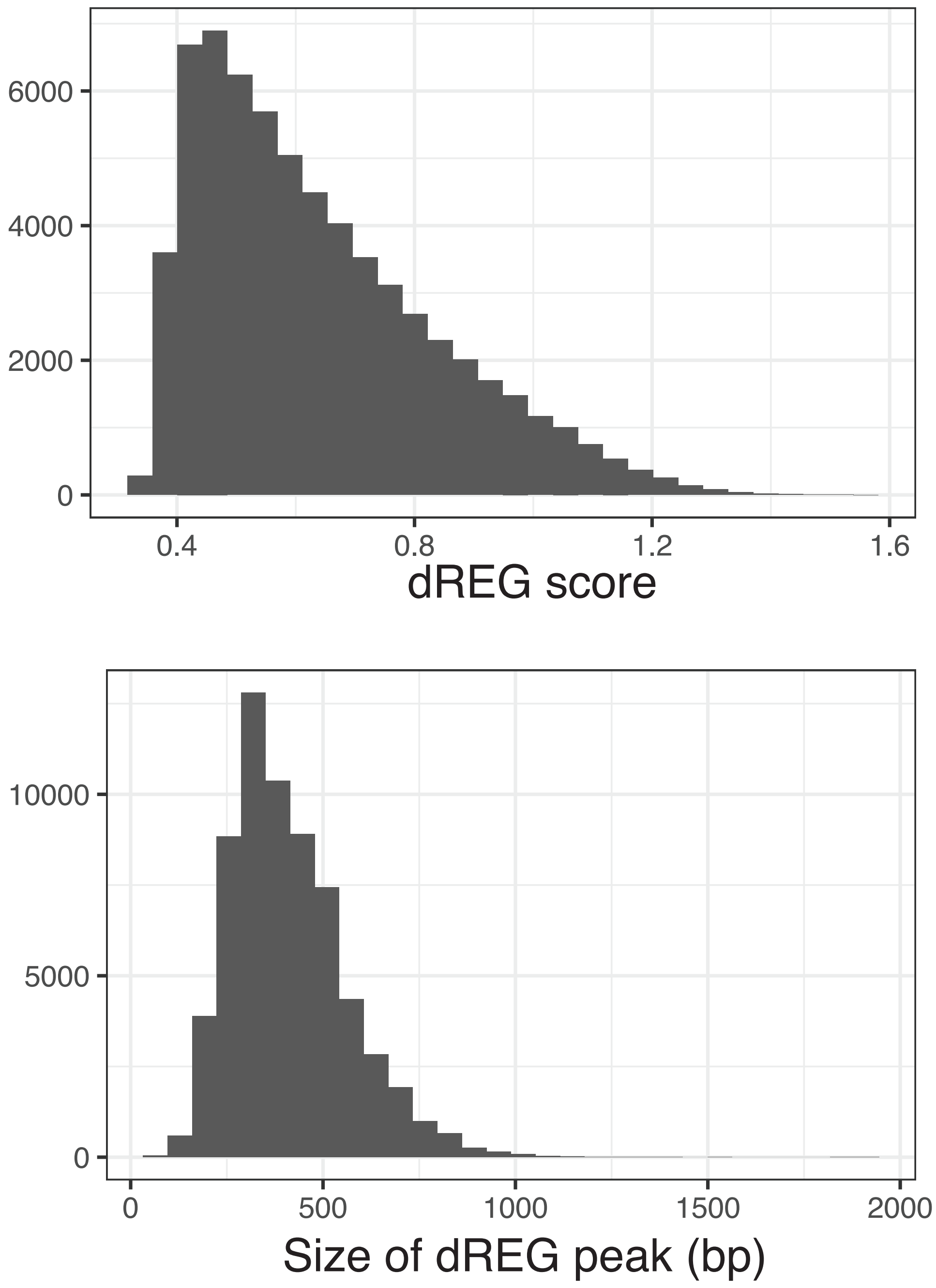


**Figure S1**. Distribution of genome-wide dREG scores (top) and dREG peak size (bottom).

Median dREG score is 0.591 and median dREG peak size is 390 bp.

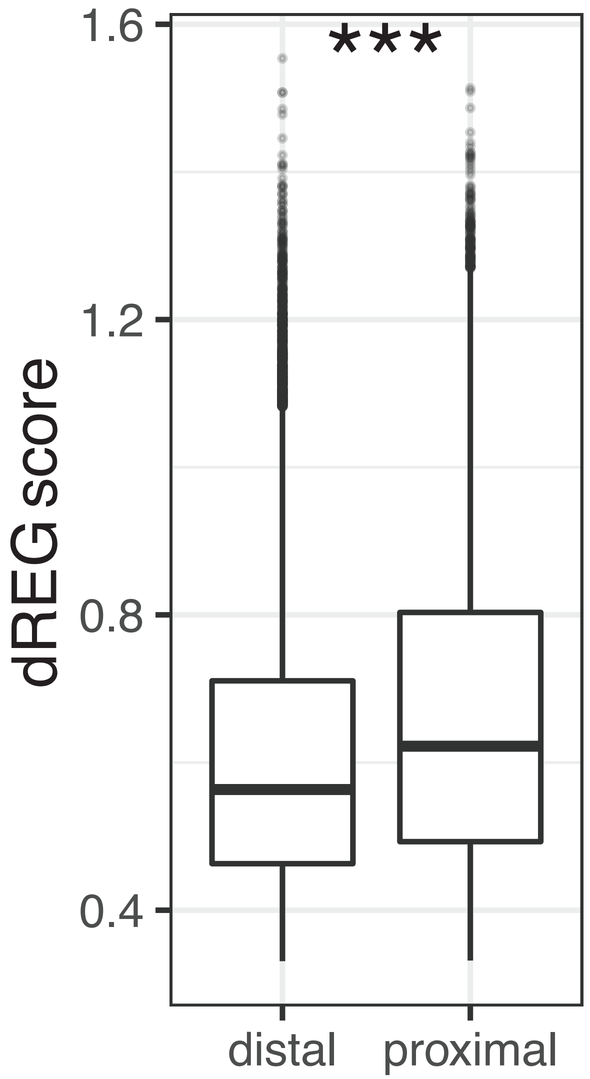


**Figure S2**. dREG scores for $dREG_{proximal}$ and $dREG_{distal}$ peaks.

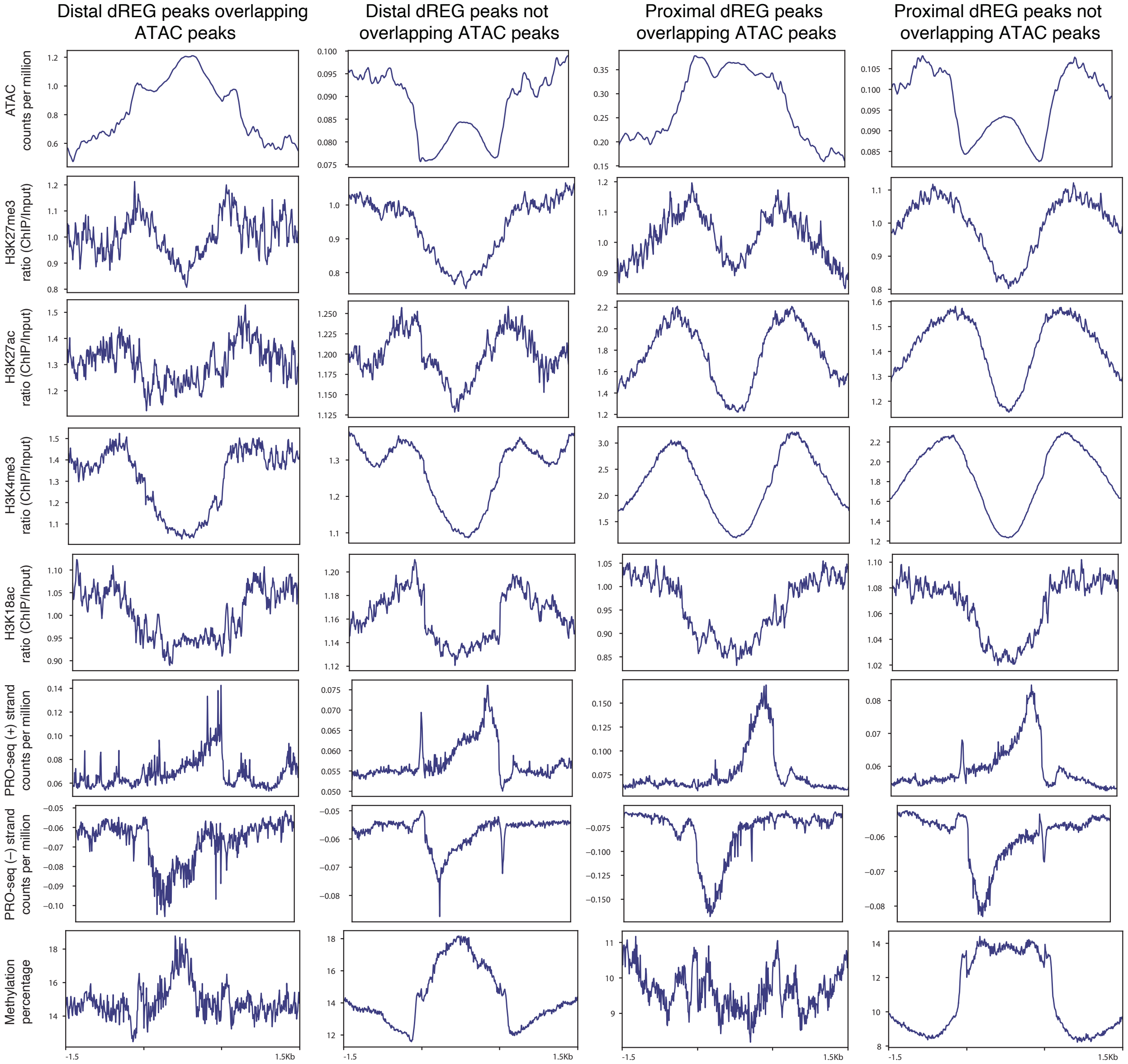


**Figure S3**. Epigenetic marks for $dREG_{proximal}$ and $dREG_{distal}$ peaks. The $dREG_{proximal}$ and $dREG_{distal}$ peaks were also divided on whether it overlapped an ATAC peak region.

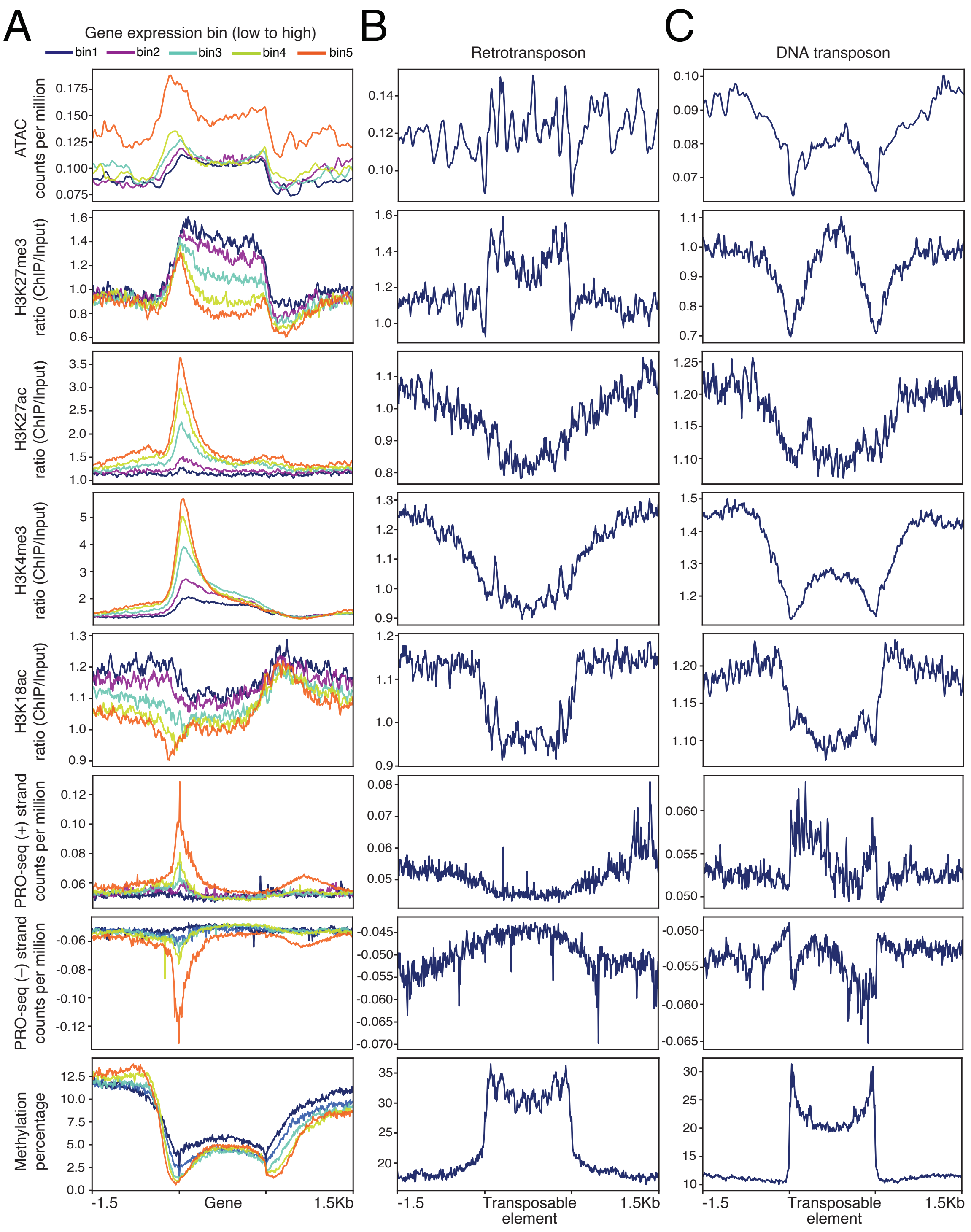


**Figure S4**. Epigenetic marks for coding and repetitive sequences in the rice genome. (**a**) Transcriptionally active genes binned by gene expression levels. (**b**) Class I retrotransposons and (**c**) class II DNA transposons.

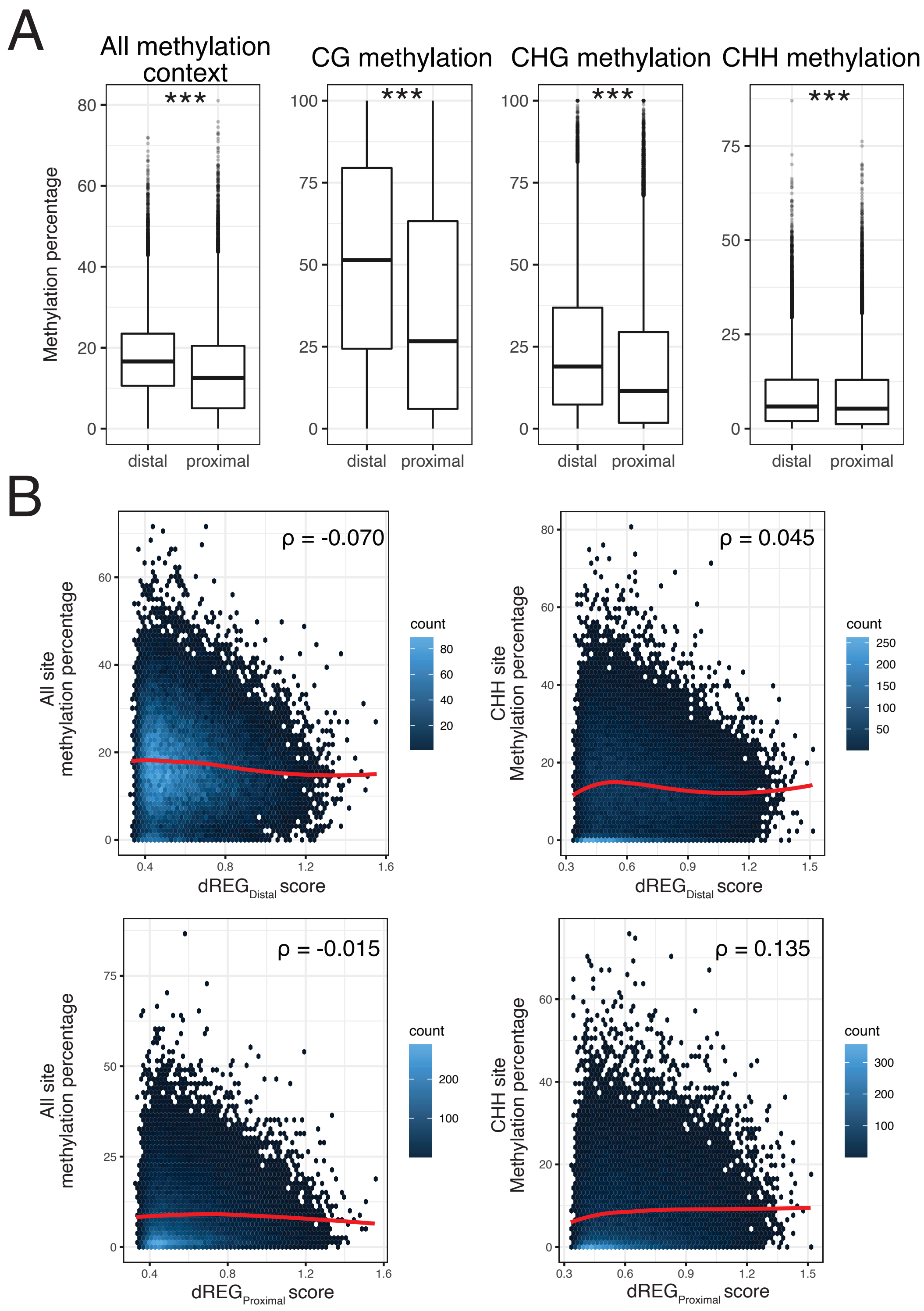


**Figure S5**. (**a**) DNA methylation levels for dREG$_{proximal}$ and dREG$_{distal}$ peaks. (**b**) Scatter plot for dREG peak regions' dREG score and DNA methylation level.

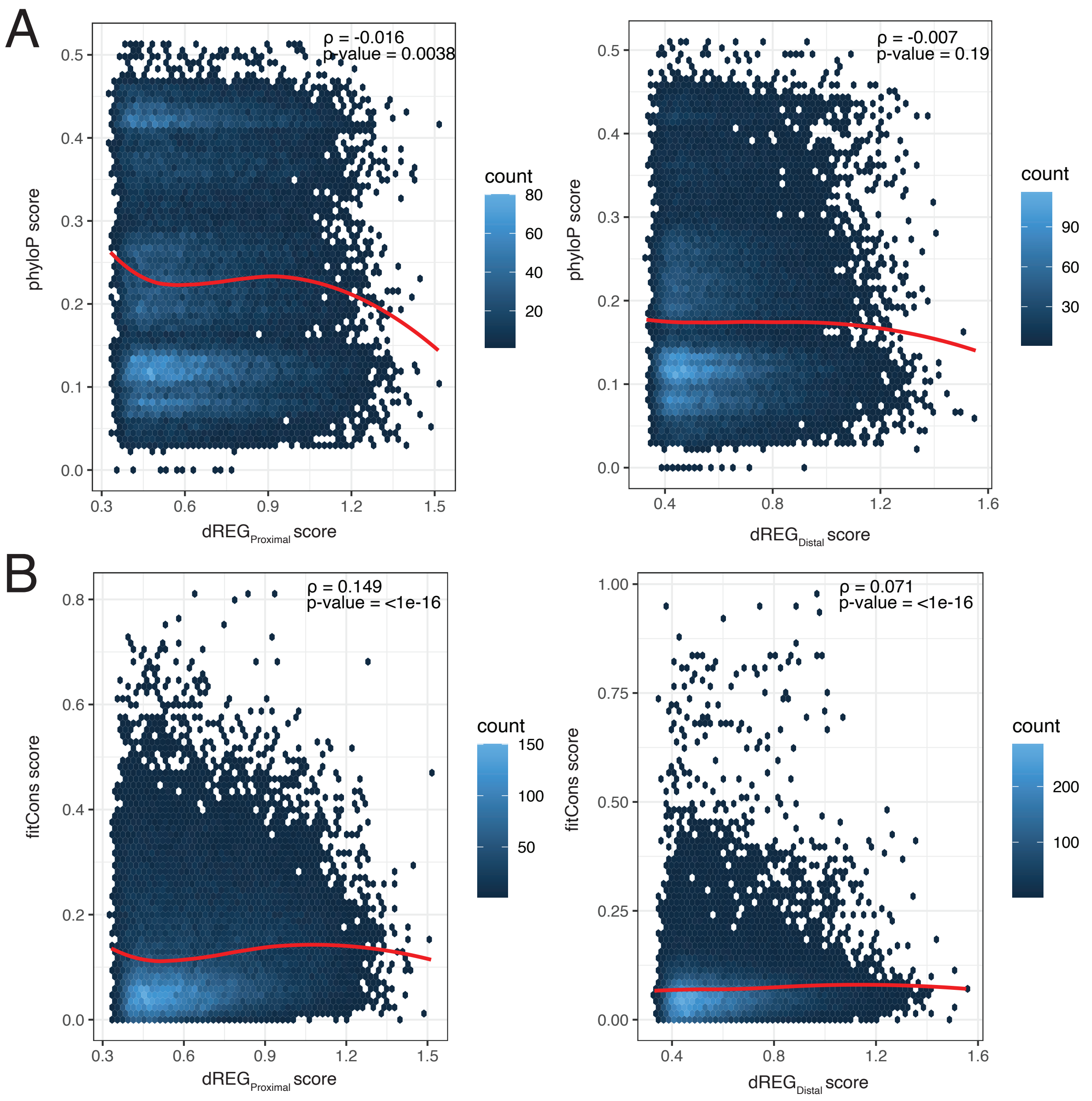


**Figure S6**. Scatter plot for dREG peak regions' score and evolutionary conservation scores. (**a**) plot for $dREG_{proximal}$ and $dREG_{distal}$ peaks with phyloP scores and (**b**) plot for $dREG_{proximal}$ and $dREG_{distal}$ peaks with fitcons scores.

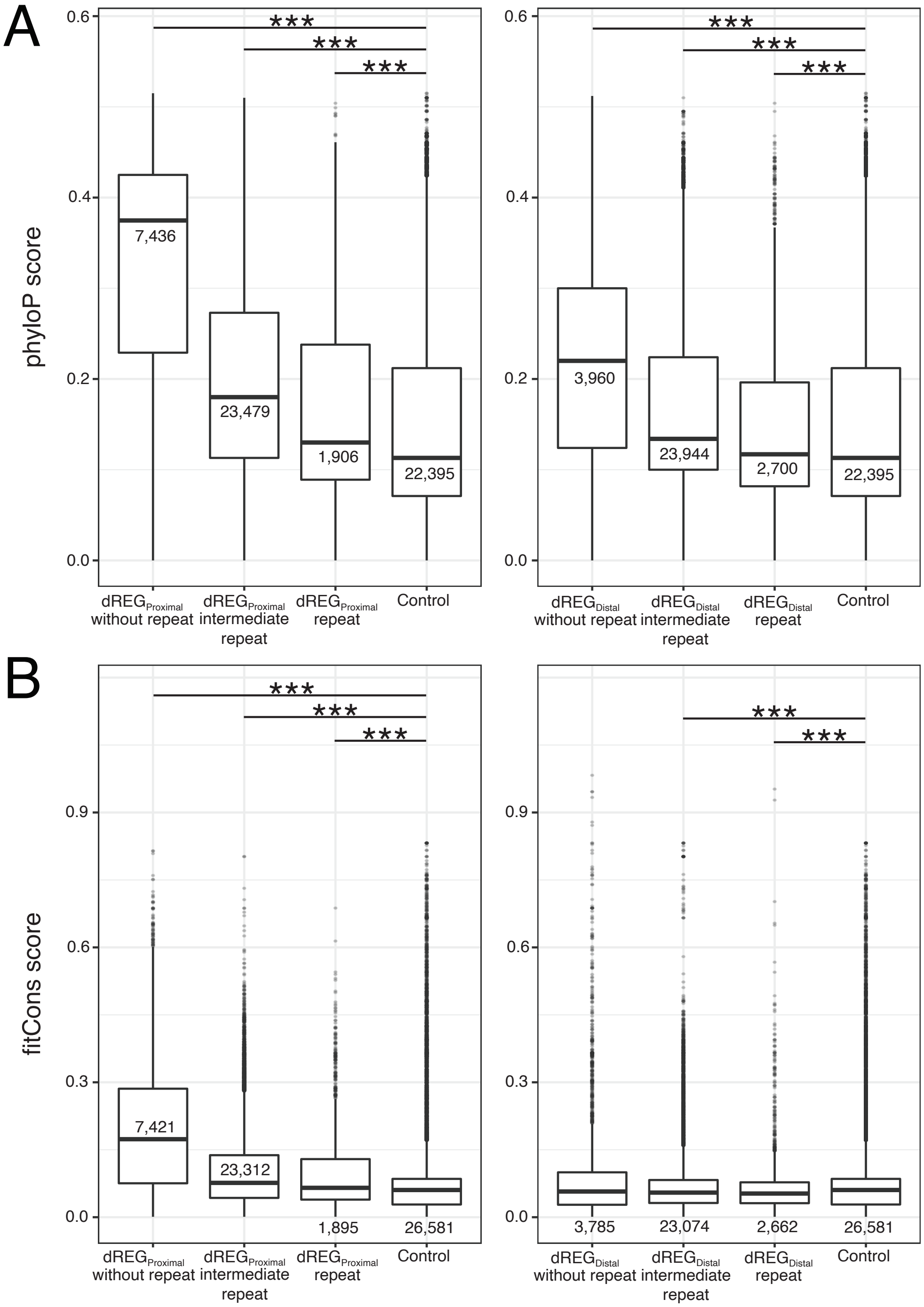


**Figure S7**. Evolutionary scores (**a**) phyloP and (**b**) fitCons for $dREG_{proximal}$ and $dREG_{distal}$ peaks. dREG peaks are divided by its repeat class. *** indicates significant differences after a Mann-Whitney test (p-value < 0.0001).

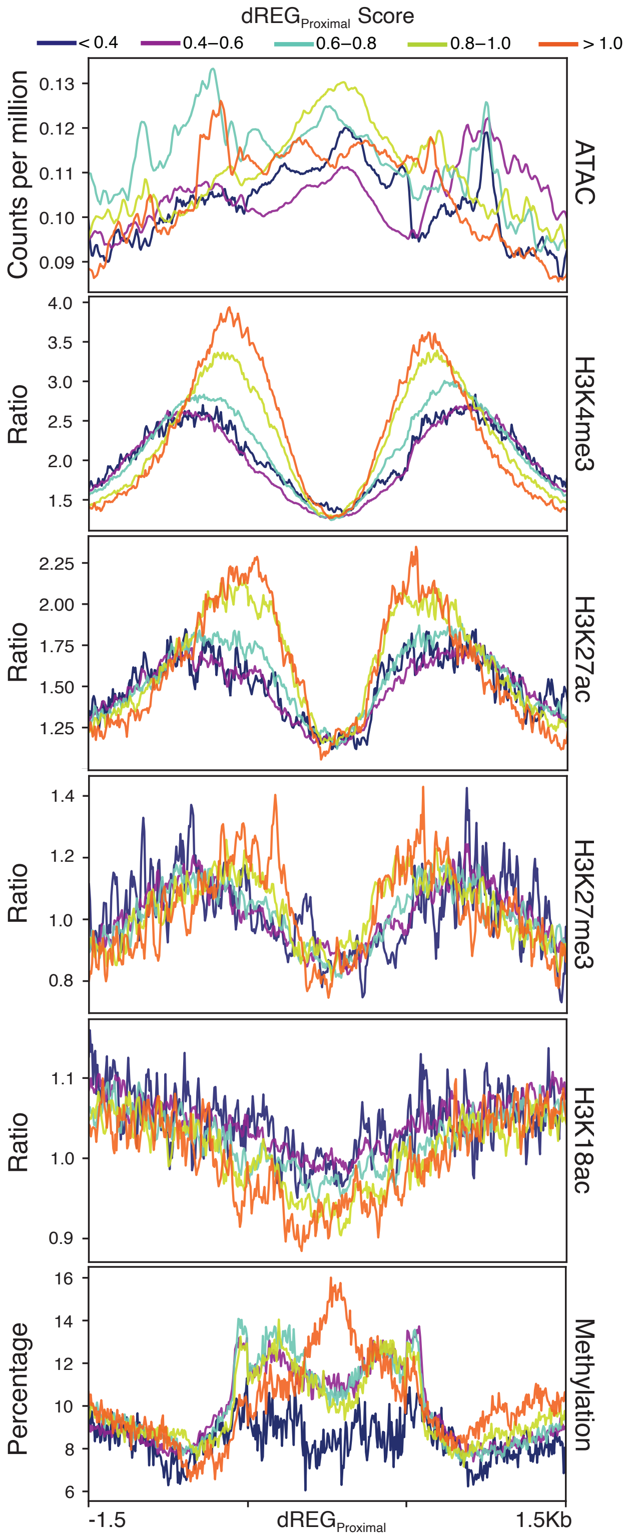


**Figure S8**. Chromatin profiles of dREG$_{Proximal}$ peaks that are binned by dREG scores.

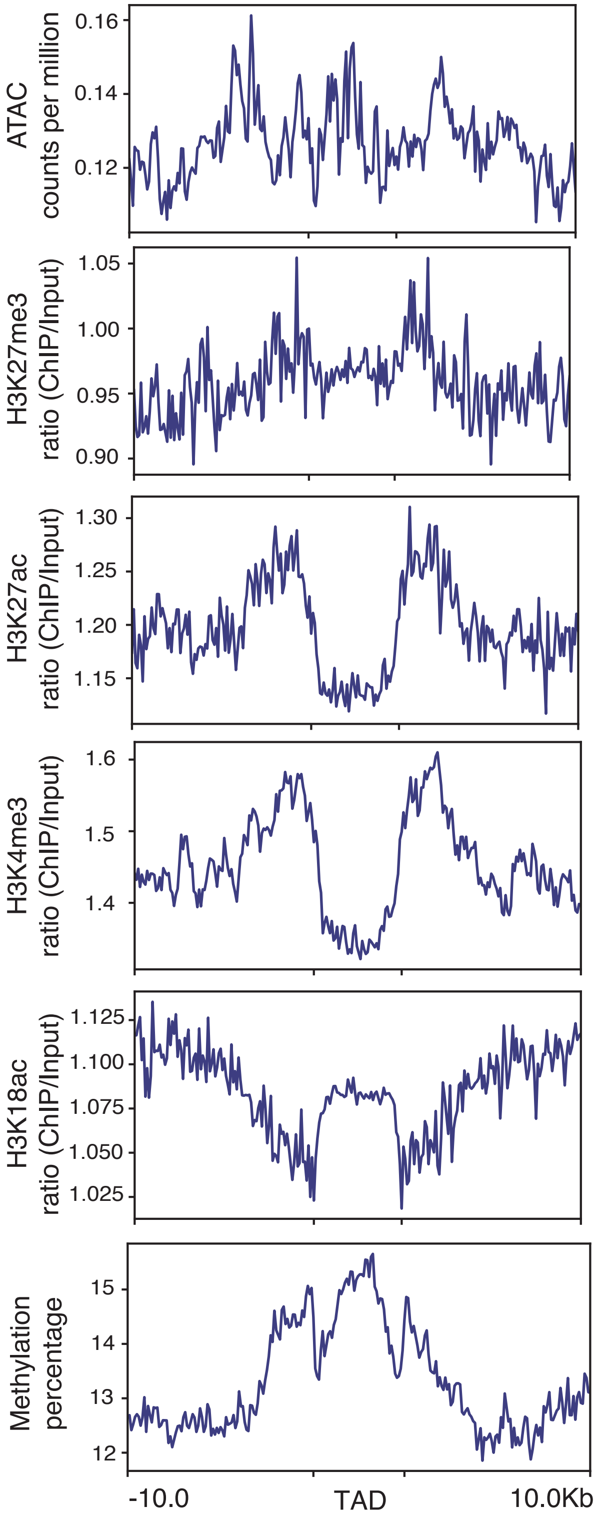


**Figure S9**. Epigenetic marks surrounding TAD boundaries.

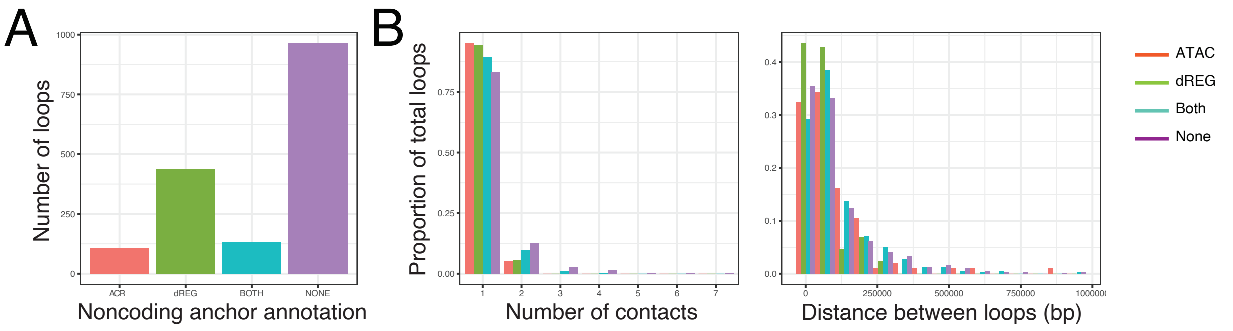


**Figure S10**. Distribution of loops detected by Pore-C for $\text{dREG}_{\text{distal}}$ peaks. (**a**) Total number of noncoding-gene loops where the noncoding anchor contains either an ATAC peak, a dREG peak, both an ATAC and a dREG peak, or none. (**b**) Across the noncoding-gene loops the number of noncoding anchor contacts per gene (left) and the distance between the gene and noncoding anchor (right).

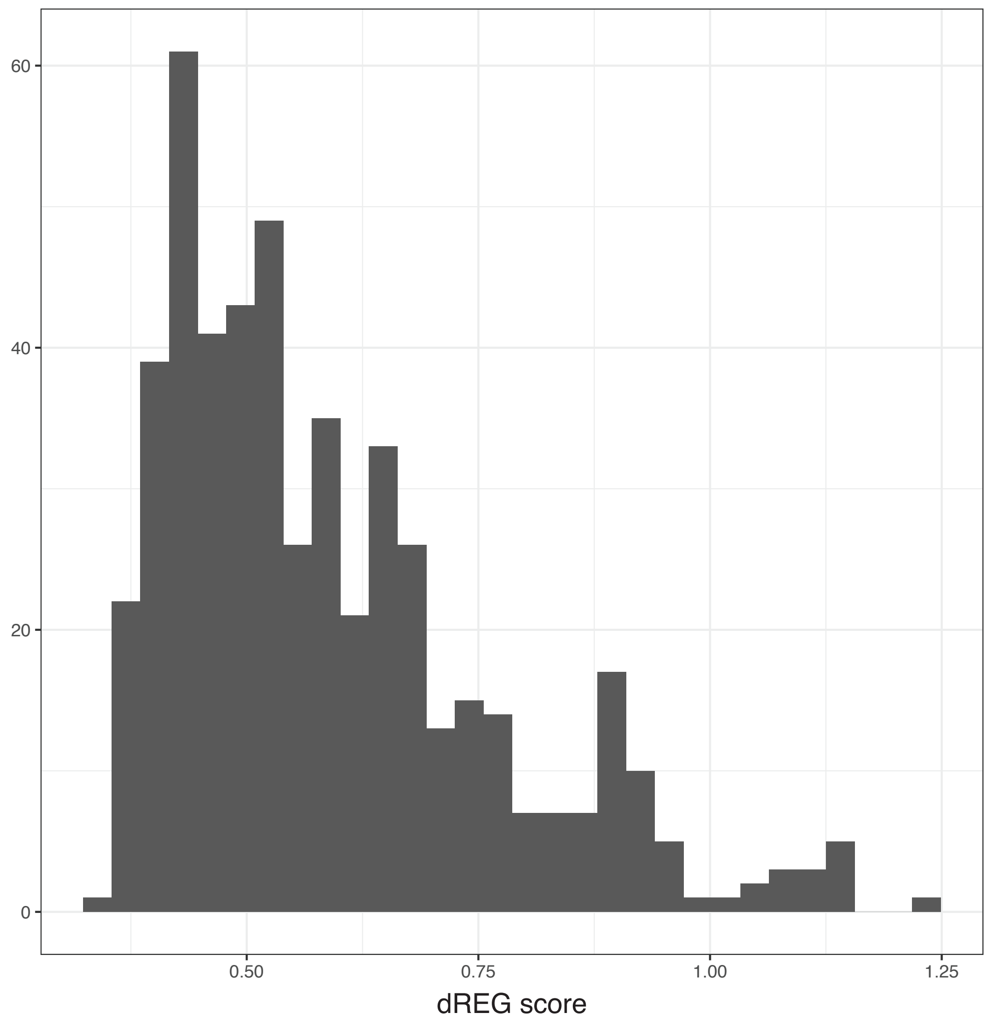


**Figure S11**. Distribution of dREG scores for the dREG peaks contacting a gene.

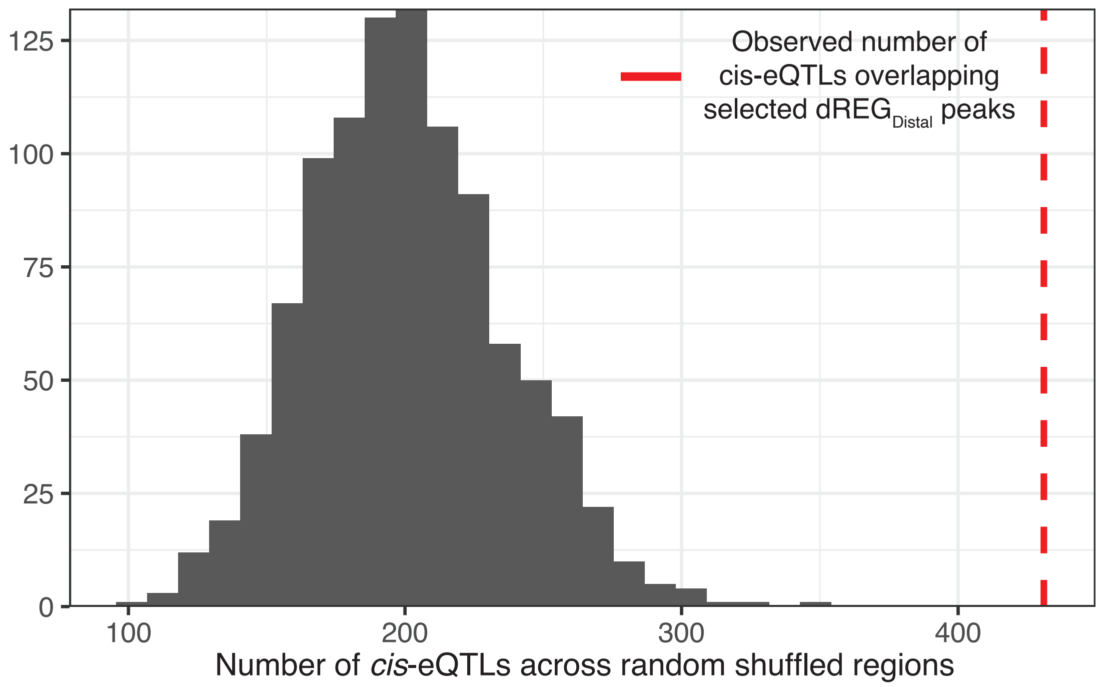


**Figure S12**. Enrichment of eQTLs within dREG$_{distal}$ peaks identified in Figure 5D

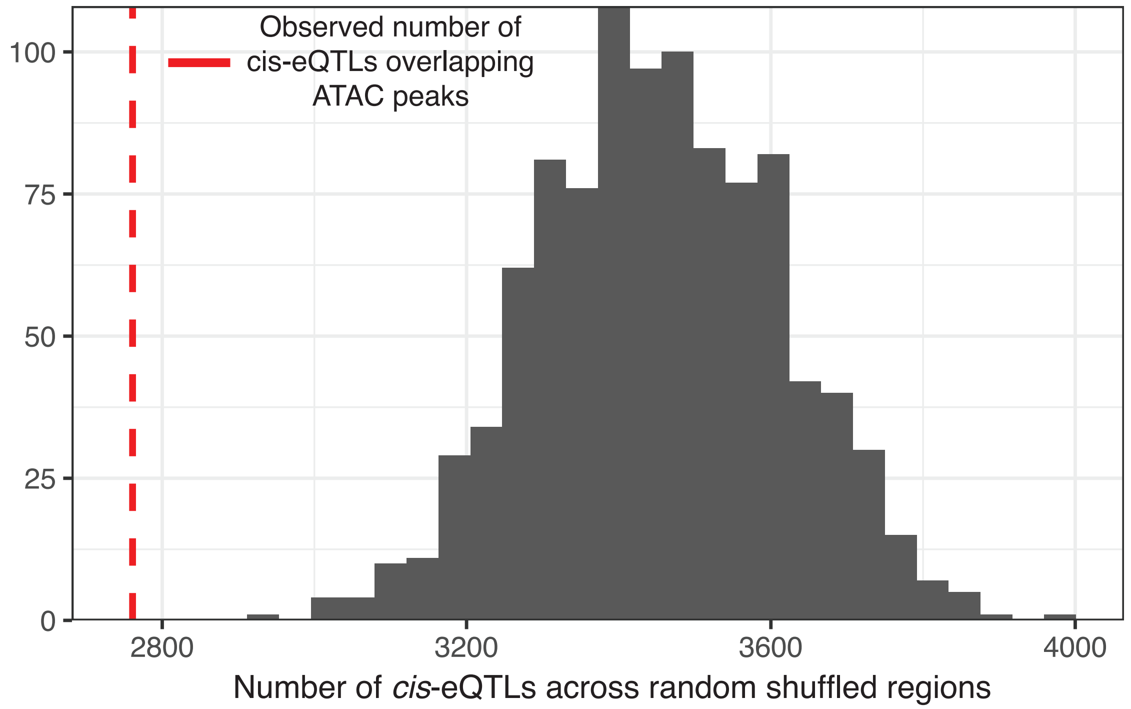


**Figure S13**. Enrichment of eQTLs within ATAC peaks.
