## Supplemental Tables for "Nascent transcription and the associated *cis*-regulatory landscape in rice"

This file includes:

Tables S1 and S2

**Table S1**. Total number and proportion of genic, distal, and proximal dREG peaks that overlapped a ATAC peak region, as well as the total number and proportion of ATAC peaks overlapping dREG peaks.

| | Not overlap ATAC | Overlap ATAC | Proportion overlap | | | Not overlap dREG | Overlap dREG | Proportion overlap |
|---|---|---|---|---|---|---|---|---|
| **Genic dREG** | 5270 | 412 | 0,0725 | | **Genic ATAC** | 2525 | 412 | 0,1403 |
| **Distal dREG** | 29540 | 1699 | 0,0544 | | **Distal ATAC** | 12997 | 1699 | 0,1156 |
| **Proximal dREG** | 29414 | 3563 | 0,1080 | | **Proximal ATAC** | 2765 | 3563 | 0,5631 |

**Table S2**. Pore-C summary statistics

| Reads | | | Number of reads with contact order | | | | | | Pairwise Contacts |
|---|---|---|---|---|---|---|---|---|---|
| Num of reads | Read bp | Contacts per Gbp | 2 | 3 | 4 | 5-6 | 7-11 | >12 | Count |
| 104 539 406 | 121 866 117 653 | 2 385 451 | 23 220 480 | 17 135 151 | 10,143,795 | 7 698 463 | 2 205 629 | 62924 | 290 705 761 |
